# Exploring Discrete Space-Time Models for Information Transfer: Analogies from Mycelial Networks to the Cosmic Web

**DOI:** 10.1101/2023.11.30.569350

**Authors:** Tommy Wood, Tuomas Sorakivi, Phil Ayres, Andrew Adamatzky

## Abstract

Fungal mycelium networks are large scale biological networks along which nutrients, metabolites flow. Recently, we discovered a rich spectrum of electrical activity in mycelium networks, including action-potential spikes and trains of spikes. Reasoning by analogy with animals and plants, where travelling patterns of electrical activity perform integrative and communicative mechanisms, we speculated that waves of electrical activity transfer information in mycelium networks. Using a new discrete space-time model with emergent radial spanning-tree topology, hypothetically comparable mycelial morphology and physically comparable information transfer, we provide physical arguments for the use of such a model, and by considering growing mycelium network by analogy with growing network of matter in the cosmic web, we develop mathematical models and theoretical concepts to characterise the parameters of the information transfer.

## 1. Introduction

The living, chemical and physical growing networks have several key similaries. First is an emergent behaviour. In all three types of growing networks, emergent behavior arises as a result of individual elements (biological cells, physical particles, chemical molecules) interacting and organizing themselves into larger structures or patterns. These emergent properties often exhibit complex and unpredictable behaviors that are not directly programmed into the individual components. Second, biological, physical, and chemical growing networks are capable of self-organization. Individual elements follow simple local rules or physical-chemical processes that collectively give rise to well-defined patterns or structures at a higher level of organization. Examples include the self-organization of cells into tissues in biological systems, the formation of snowflakes in physical systems, and the crystal growth in chemical systems. Third, growing networks in all three domains exhibit a degree of adaptability and flexibility. Biological networks, such as neural networks, can rewire connections and modify their structure in response to environmental changes. Physical networks, like fractals, can adapt their geometric patterns depending on growth conditions. Similarly, chemical networks can adjust reaction rates and equilibrium positions based on external factors. Fourth, all growing networks tend to optimize certain properties. In biological systems, evolutionary processes favor the development of efficient and adaptive organisms. Physical networks often evolve towards efficient and stable structures due to physical principles like minimum energy states. In chemical systems, the reaction pathways can optimize to achieve the most stable and energetically favorable products. Fifth, growing networks often exhibit hierarchical organization, where structures are composed of smaller subunits, which, in turn, can be composed of even smaller components. This hierarchical arrangement is seen in biological systems (e.g., tissues composed of cells), physical systems (e.g., branching patterns in rivers or trees), and chemical systems (e.g., complex molecules made of simpler building blocks). Sixth, feedback loops play a crucial role in all growing networks, ensuring the stability and regulation of the system. In biological systems, feedback mechanisms control processes like cell division and differentiation. Physical systems often involve feedback loops that influence the growth and branching patterns. In chemical networks, feedback mechanisms can lead to autocatalytic reactions, where a product enhances its production. Seven, growing networks often operate in non-linear and non-equilibrium regimes. In biological systems, processes like morphogenesis or tissue formation involve non-linear interactions between genes and proteins. Physical systems, such as turbulent flow, are also characterized by non-linear dynamics. Chemical reactions often occur far from equilibrium, leading to dynamic behaviors like oscillations and pattern formation. In present paper we develop a formal theory of information propagation in growing mycelium networks by analogy with space matter networks.

In [1] we proposed that slime mould *Physarum polycephalum* growing on a nutrient agar might show some analogies with an expanding universe: high density of the slime mould body at the front of the propagating wave being transformed into a network of protoplasmic tubes further from the proximity. Two illustrations of our experiments are shown in Fig. 1. There is evidence, e.g. as illustrated in Fig. 1a, of the slime mould growth being similar to excitation wave front [2, 3, 4]: an excitation part corresponds to actively growing mycelium and zone of fruiting body, while a refractory part relates to a substrate exhausted of nutrients.

**Figure 1.**
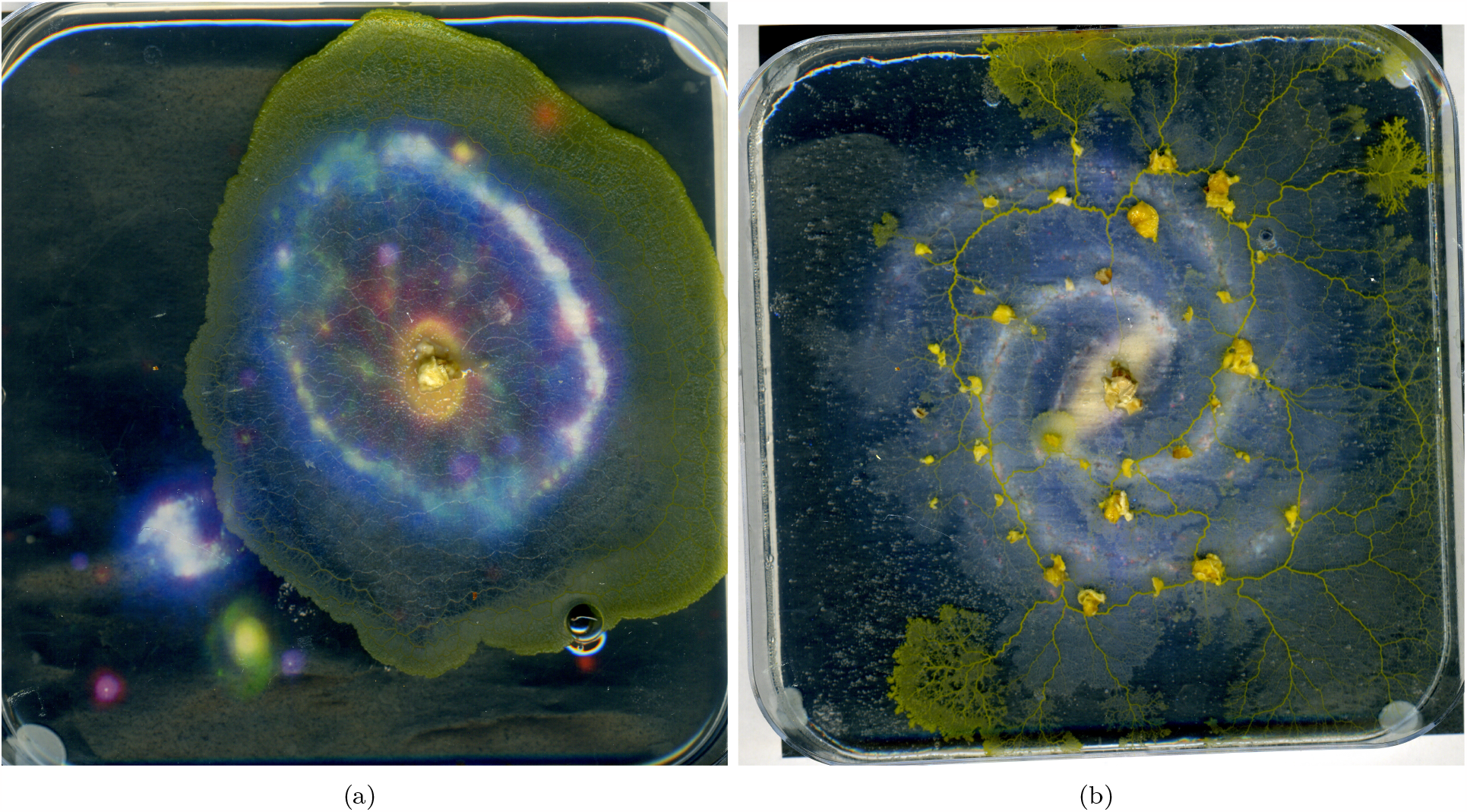
Slime mould *Physarum polycephalum* growing on a template of the (a) Cartwheel Galaxy and (b) Galaxy. From [1].

In the form of slime mould, fungi propagate underground as waves of mycelium growth. The rings were first described in detail in 1917 in relation to fairy rings in eastern Colorado [5]. The paper [5] demonsrated that mycelium wave fronts can be detected by richer and denser vegetation. In modern days the rings can be found using Google maps, as illustrated in our personal example in (Fig. 2ab). Thus, when thinking about large-scale fungal waves we can see the following scheme, see Fig. 2c, a propagating wavefront, which can also show fruit bodies, and a tail of mycelium network.

**Figure 2.**
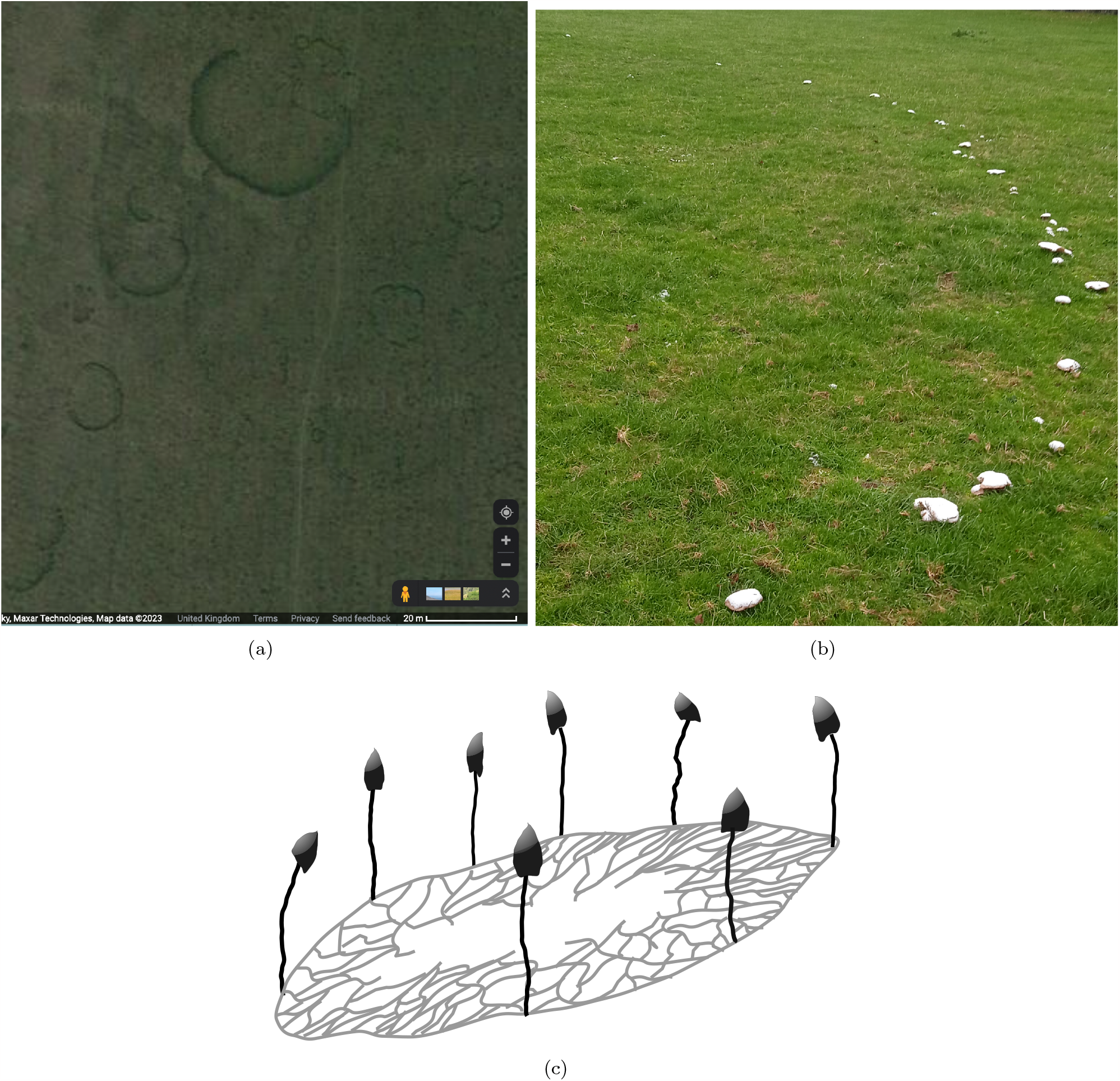
Fairy ring, wave-fronts of propagating mycelium of the parasol mushroom *Macrolepiota procera* spotted using Google maps (a) and verified by site visit (b). Location of the site in North Somerset, near Bristol, UK, exact coordinates are available on request. Photo (b) is by Andrew Adamatzky. (c) A scheme of a propagating fungal galaxy

New computational methods for modelling matter distribution between galaxies inspired by *P. polycephalum* [6] have been shown to not only corroborate observed intergalactic structure, but also to imply some yet unknown connection between emergent biological forms, large scale cosmological structure and, as will be explored in this paper, discrete microscopic space-time.

Significant difficulties involved with mathematically unifying general relativity and quantum field theory have been the primary focus of theoretical physicists over recent decades [7]^1^. This has led to a number of attempts at unification such as string theory [8, 9, 10] for its prediction of an interacting graviton, as well as loop quantum gravity [11, 12, 13, 14, 15] as a mechanism for renormalizing gauge theories of gravity.

These theories, despite being incredibly successful at producing mathematical and conceptual connections still fail to generate a complete solution [16] and still have not yet been able to be varified by experimental observations of theoretical predictions [17]. Gravity invovles space-time geometry and quantization of gravity naturally leads to the concept of quantizing spacetime, implying atoms of space and atoms of time [18, 19]. The following questions then arise. How are the atoms of space arranged? How does this arrangement evolve through discrete time? How can such a particular arrangment lead to emergent consciousness and at the correct scales, the observed behaviour of our universe? Computers and data structures provide researchers with indispensible tools with which to probe such questions and in recent years have led to a number of theories based on connections between atoms of space, these include, cellular automata or networks [20, 21, 22, 23, 24, 25, 26, 27, 28, 29, 30, 19, 31], causal sets [32, 33, 34, 35, 36, 37], and causal dynamical triangulations [38, 39, 40, 41], each treating spacetime as constituted of indivisible discrete units of space.

More recent work by Gorard and others has demonstrated that such discrete networks, with specific substitution rules, are not only numerous and compatible with physical laws [42, 43, 44, 45, 46, 47], but have a whole host of applications relating to the symbolic structure of mathematics, logic and algorithms, implying underlying structural connections between of what could be the foundations the structure of space-time and relations in abstract areas of thought [48]. There is some demonstration of the compatibility with special and general relativity and a structure for understanding quantum mechanical properties. However as stated, the number of potential discrete universe-generating network update rules are numerous and homing in on one specific structure is an open problem. A discrete space-time description must have the capacity to regenerate all observed physical laws without contradiction.

One unifying theme with the discrete spacetime theories proposed is that of emergence, that the complexity and chaos observed in nature comes out of underlying simple structure and rules. A good example of this is the game of life [23], where simple underlying rules generate complex behaviour. Emergence is discussed in almost all disciplines [49], from emergent discrete spacetimes [27, 50, 33, 51, 30], to the idea of gravity being an emergent phenomena, [52, 53], to biological systems [54, 55, 49], language [56], computer programmes [26], and consciousness [57, 58], to name but a few. It is a powerful and very fundamental idea that can be seen as taking effect on a variety of different scales.

The *Space Element Reduction Duplication (SERD) network model* is a discrete spacetime model developed in [59, 60, 61], the fundamental rules and structure of which emerge from fundamental philosophical principles. Out of these fundamental principles emerge rules, which themselves give rise to physical comparisons and biological dendritic radial topology^23^. This makes this system an ideal system for probing the apparent connection between biological and large scale cosmic structure, and its possible relation to underlying space-time topology [6].

In this article we explore the connections between emergent network structures in early life forms such as *P. polycephalum* [54] and *mycorrhizal fungi* [62] and the cosmic web and hypothesise on possible space-time topologies based on these structures, in particular the SERD network.

The cosmic web is a network of matter and dark matter that fill the spaces between luminescent galaxies [63, 64] and has been shown to follow the Voronoi diagram structure generated by *P. polycephalum* [65], an idea that has only been strengthened by insights from [6].

Section two gives a brief overview of our understanding of the science of mycelium networks. Section three introduces the structure and rules of the information transfer network of the SERD model and provides motivation for its use as both a cosmological model and a model for mycelium growth and information transfer. In Section four we show some topological correspondences between the emergent SERD network structure and early life structures such as mycelium networks Fig. 24 [66, 67, 68, 65, 69], and, by using an information-energy correspondence, we demonstrate its compatibity with cosmological observations of the redshift scale factor relation implied by the FLRW metric from the ΛCDM model [70, 71, 72, 73].

The SERD network generates a mycelium-like hypergraph structure that evolves as a multiway system through time and incorporates information propagation as an emergent property. We consider how information propagates in this system, and make connections with the manner in which information travels within early evolved living systems and the cosmic web. Perhaps the naturally occurring networks in nature will provide us with clues as to what the underlying network structure of space-time may be.

## 2. Science of Mycelium

Anywhere on land that is not a desert, bear rock or an over-developed urban environment, contains an expansive network of connected mycelium just below the surface. Mycelia are one of the oldest and widely distributed groups of organsms on earth [74].

Mycelia or mycelial fungi^4^, are the root-like networks of connected *hyphae* of the fungal organism. The *hyphae* are thread-like tubes of consecutive cells, separated by *septa*, that allow for the controlled flow of organelles and nutrients, as well as cytoplasmic continuity [75, 76].

Starting from a single spore^5^ a single germ tube^6^ extrudes and undergoes a period of apical extension, this is followed by apical and sub-apical branching^7^. This network then expands, growing equally in all directions as a *fractal spanning tree* [66, 77], *with growth concentrated at the apex as the network spans out in search for nutrients. Fungi must forage for resources, normally dead organic matter, and use its mycelial network to transport nutrients between source and sink regions, thus the importance of porus septa* and continuity between cells [75].

As these networks grow via hyphal tip growth and branching, a form of hyphal fusion may occur known as *anastamosis*, where the walls of two adjacent hyphae fuse allowing a junction to form between them. This can increase the connectivity of the mycelial networks allowing for greater resilience to breakages in the network caused by cell death, due to grazers or lack of resources [78], and may shorten the shortest distance for nutrients to travel between source and sink^8^.

When the mycelial fungi find an abundant nutrient source, highly conductive channels known as *chords* form between source and the foraging fronts. Chords are normally produced when neighbouring aligned hyphae aggregate, creating a thicker and more insulated path for nutrients to flow through. This tends to increase flow rate of nutrients between a source and sink.

Hyphae that are in nutrient-depleted regions with no usefulness for transport form a *regression* region. These regions show a decrease in biomass of the mycelium as organelles, nutrients and cellular contents are recycled and transported for use in other parts of the network [75].

When resources are sufficient and conditions favorable the mycelia of the fungal organism will, via coordinated hyphal growth, grow spore-producing fruiting bodies. These will spread the organsims genetics and allow it to reproduce. The new spore will, via dispersion, find a new location and then act as a new hyphal hub. A mycelial network may have many hyphal hubs originating from a number of spores.

This process of hyphal growth, branching, fusion and regression results in mycelium being highly adaptable to its environment, growing new branches in search for food, recycling its own biomass [66] and forming channels of varying conductivity.

As well as facilitating the flow of nutrients and organelles, hyphae also act as channels for waves of electrical potential [79, 80, 81, 82, 83]. Recent observations into the electrical spiking of the fungal organism show, links between spiking patterns and environmental stimuli[84, 85]. A spike varies in its duration from 1 to 21 h [86]. This implies slow processing^9^, however fungal computation has been shown to be capable of nontrivial mappings of electrical signals. This is largely influenced by the novel electrical characteristics of the substrate as well as unique network topology and morphology [87]. They perform decentralized multi-parallel computational tasks, while having morphological adaptability^10^.

The hyphae act as bi-directional channels for both cytoplasm and electric current. Each cell in the hyphae are separated by *septa* that contain pores that allow for the flow of cytoplasm and organelles and make it easier for electrical potential to be transferred between cells. These septa may become blocked by organelles called *Woronin bodies* when the mycelium is damaged. These have the effect of blocking the pores, impeding the cytoplasmic flow. They also play a role in impeding the flow of electrical information along the hyphae, effecting which regions of the mycelium are effected by electical signalling [88]. These electrical signals travel far, transferring information between fruiting bodies of a cluster of fruit bodies[85].

Mycelium is one of the earliest multicellular organic structures [74] and therefore is of broad scientific interest, from the emergence of the morphology of the earliest multicellular organisms to underlying physical laws and how they may govern the emergence of this specific morphology [89]^11^. In this paper we propose the hypothesis that currents of both energy and matter follow along hyphae-like threads of atomized space, that space itself, at the most fundamental of levels, has a similar morphology to that of early life and so in order to provide greater access to its currents, life naturally evolves in a manner which follows a trace of the energy flows of the underlying stucture of space-time.

## 3. Modelling Methods

### 3.1 Space Element Reduction Duplication (SERD) network overview: structure and rules

The SERD network is an emergent highly dynamic background independent space-time model with bidirectional propagating and interacting wave packets of topological information. It is constituted of two fundamental elements, the *hyperedges* and the *edges*, with the boundaries between elements being the *nodes* of the network. The elements are the *Point Particle* (PP) (the hyperedges)^12^ and the *Space Element* (SE) (the edges)^13^, while the nodes exist between the elements and are refered to as *Information Gaps* (IGs), and may store the propagating topological information of the SERD network^14^. More up-to-date details on this model may be found at [61], however an overview will be provided here for convenience.

#### 3.1.1 Formal definitions for computational implementation

The topology of a given state of the SERD network is defined by the elements in the state. This information is stored in separate sets, the ***Point Particle set*** and the ***Space Element set***. An ***Information Gap set*** is also included, which allows access to the information of neighbouring elements to IGs that is useful for information propagation.

IGs exist between all neighbouring elements, they are the nodes of the network and store the topological information that propagates through it. Each IG in the network is unique and defined by a given string. In computation this may be *x*1 or *x*5262 for example^15^.

##### Definition 3.1.

*A* ***Space Element set*** Σ(*t*) *is an edge set that stores the information of all SEs that exist in the state of the SERD network at a given time t*.

##### Definition 3.2.

*A* ***Space Element index set*** Σ_*in*_(*t*) = (*n*_1_, …, *n* _|*S*(*t*) |_), *stores the addresses of each SE within the SE set. n*_*i*_ *maybe either a natural number or a string depending on whether* ***arrays*** *or* ***dictionaries*** *are used for storage of the SE sets*.

**Table 1:**
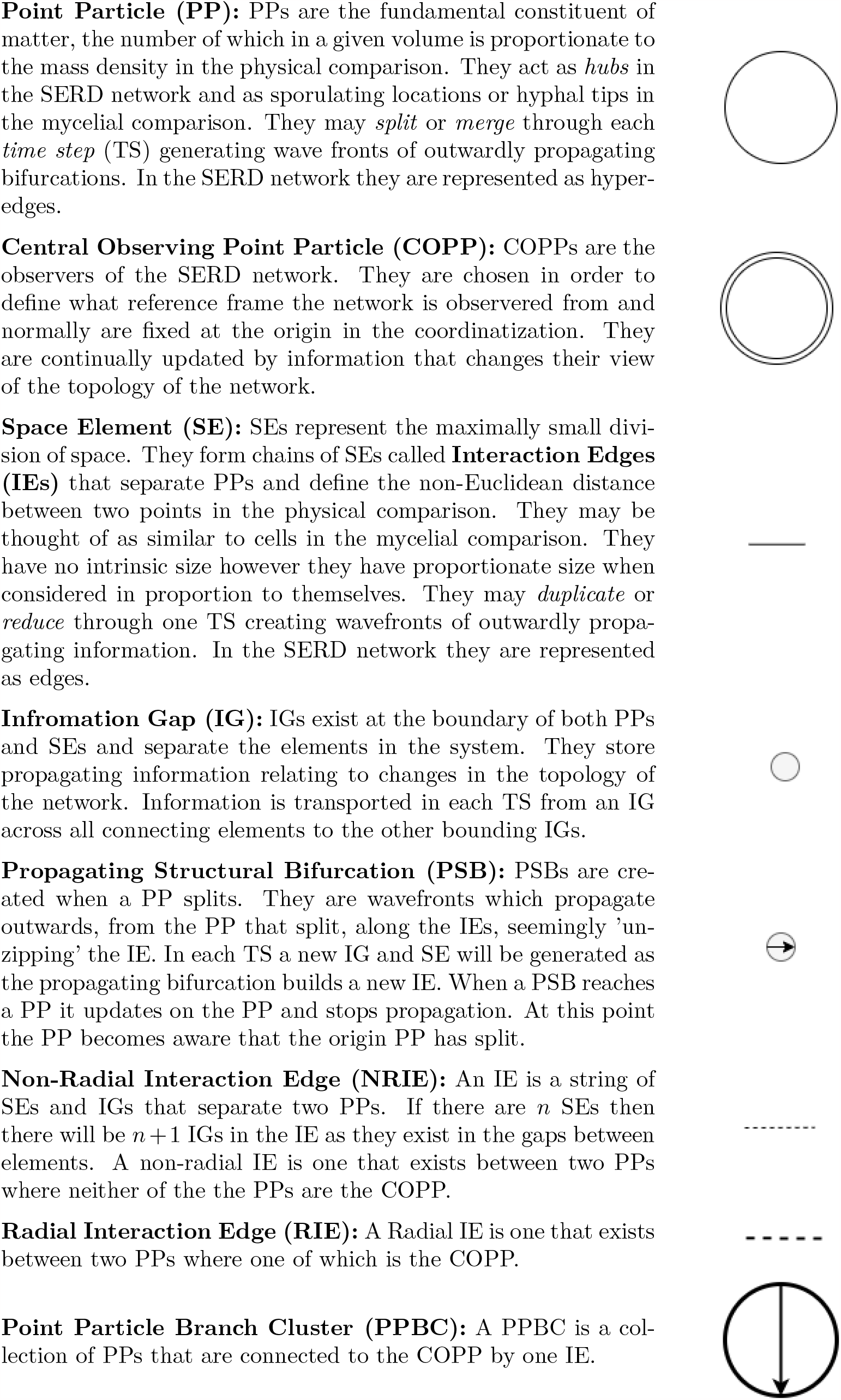
Symbolic diagram key.

##### Definition 3.3.

*A* ***Space Element*** *s* = (*′xn*′,′ *xm*′) *with s* ∈ Σ(*t*) *is an edge that connects two IGs, xn and xm, in the SERD network. SEs are labelled by an alphanumeric string starting with the letter s concatenated with a natural number n*.

##### Definition 3.4.

*A* ***Point Particle set*** *P* (*t*) *is a hyper-edge set that stores the information of all PPs that exist in the state of the SERD network at a given time t*.

##### Definition 3.5.

*A* ***Point Particle index set*** *P*_*in*_(*t*) = (*n*_1_, …, *n* _|*P* (*t*) |_), *stores the addresses of each PP within the PP set. n*_*i*_ *maybe either a natural number or a string depending on whether* ***arrays*** *or* ***dictionaries*** *are used for storage of the PP set*.

##### Definition 3.6.

*A* ***Point Particle*** *p* = (*′xn*′, …,′ *xm*′) *is a hyper-edge containing an arbitrary number of IG labels, storing information of all neighbouring IGs. PPs are labelled by an alphanumeric string starting with the letter p concatenated with a natural number n*.

##### Definition 3.7.

*An* ***Information Gap set*** *X*(*t*) *is a set that stores the information of all IGs which exist in the SERD network at a given time t*.

##### Definition 3.8.

*An* ***Information Gap*** *is a mathematical object that stores information of surrounding elements, PSBs hosted on the IG and any topological information propagating through the network. It can be written as x* = ((*′pq*′, …,′ *pr*′), (*′sn*′, …,′ *sm*′), …*I*…), *where x* ∈ *X*(*t*) *and I represents an arbitrary object of propagating information*^16^. *IGs are labelled by an alphanumeric string starting with the letter x concatenated with a natural number n*.

#### 3.1.2 Space Element (SE) actions

As the SERD network evolves through discrete time it’s elements may or may not perform an *action*. This action, for an SE, may be either to *duplicate* or to *reduce* as represented in Fig. 3. These actions are thought to occur with the same weighting, however it is possible for output states to have incurred consistant imbalance in the number of reductions and duplications occurring and so are of particular interest, when considering things like information density in an expanding universe.

**Figure 3.**
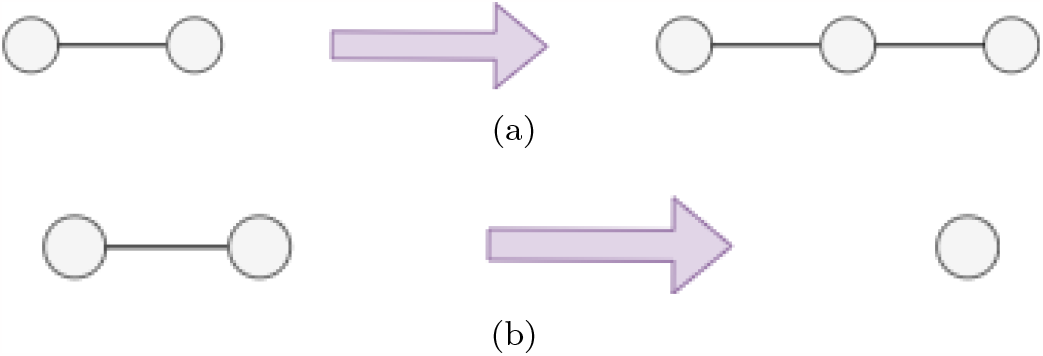
A simple diagramatic representation of the *duplication* (a) and *reduction* (b) respectively of one SE bounded by two IG nodes. In (a) information is unchanged on the boundary and an empty IG inserted. In (b) information is superposed.

#### Duplication

*One SE spontaneously becomes two SEs through a TS. A bit of information relating to the action is then created at the bounding IGs. Information is preserved at boundary IGs. A new empty IG is created between the two new SEs*. (…, *s* = (*′xn*′,′ *xm*′), …) → (…, *s*′ = (*′xn*′,′ *xq*′), *s*^*′′*^ = (*′xq*′,′ *xm*′), …) *where s, s*′ *and s*^*′′*^ ∈ Σ(*t*) *and* (…, *x, y*, …) → (…, *x, y, z*, …) *where x, y, z* ∈ *X*(*t*) *and y* = ((), (*s*′, *s*^*′′*^), …()…) *where* () *represents an empty set*.

##### Reduction

*One SE spontaneously disappears*, (…, (*′xn*′,′ *xm*′), …) → (…, …). *This causes the information stored on the two bounding IGs to superpose* (…, *x, y*, …) → (…, *x* ⊕ *y*, …) *where x, y* ∈ *X*(*t*) *and the operation* ⊕ *represents the sharing of information (neighbouring elements and propagating information) between the two IGs onto the remaining IG*^17^.

###### Definition 3.9.

*The* ***effective probability of reduction*** *is denoted by p*_*r*_ *and may be calculated in a time interval by the equation* 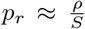, *where ρ represents the total number of reductions that occur in a time interval and S represents the total number of SEs that may or may not reduce through that time interval*.

###### Definition 3.10.

*The* ***effective probability of duplication*** *is denoted by p*_*d*_ *and may be calculated in a time interval by the equation* 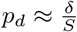, *where δ represents the total number of duplications that occur in a time interval and S represents the total number of SEs that may or may not duplicate through that time interval*.

#### 3.1.3 Point Particle (PP) actions

As a PP evolves from one TS to the next it may perform a *split* as represented by Fig. 4. If it is connected to another PP separated by a single IG it may *merge* however this action is not included in the investigation in this paper.

**Figure 4.**
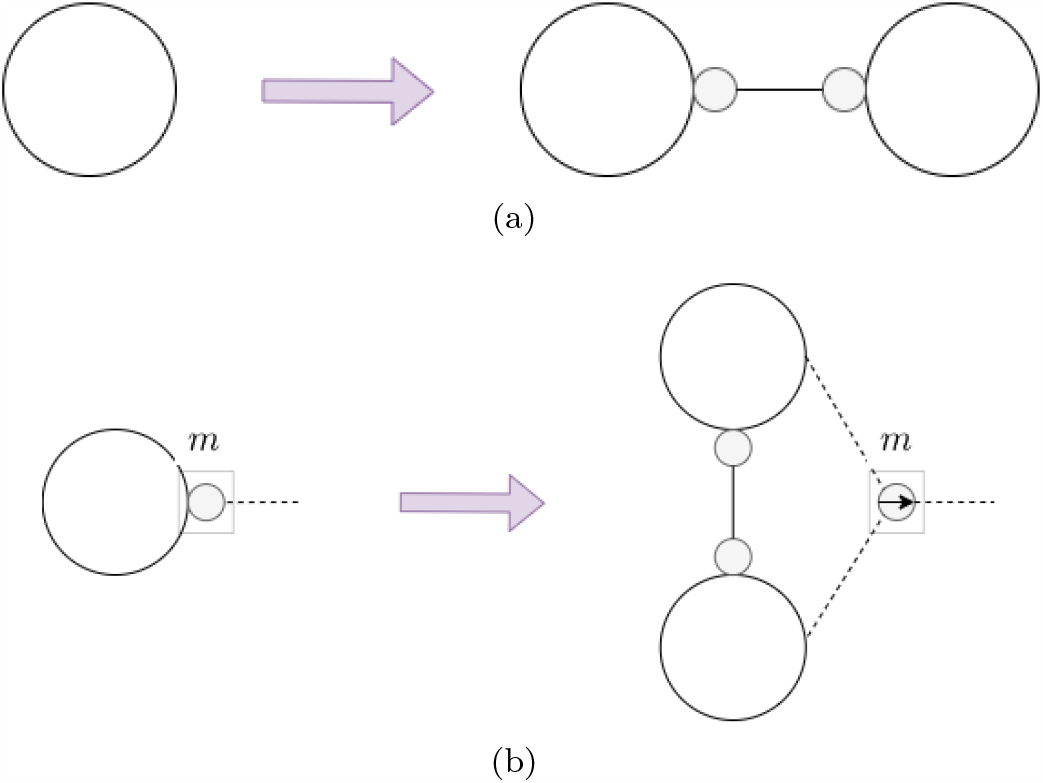
Diagrams showing the update operation of a split - both at the first instance in time (a) and a general split at a moment in time after that (b). A new IE is created of length 1 SE between the two new PPs. *m* ∈ ℕ is the number of IGs bounding the PP.

##### Split

*When one PP spontaneously splits in two through a TS. This produces two PPs from one. All connections on the produced PPs from the original are conserved*. (…, *p* = (*′xn*′, …,′ *xm*′), …) → (…, *p*′ = (*′xn*′, …,′ *xm*′), *p*^*′′*^ = (*′xn*′, …,′ *xm*′), …). *Propagating Structural Bifurcations (PSBs) are created on all neighbouring IGs. These transport the information of the split along the IEs to all other PPs in the system*.

##### Merge

*A merge is where two PP merge or fuse together to become one. This may involve a sharing, or superposition of neighbouring IGs between the two merging PPs*^18^.

###### Definition 3.11.

The *effective probability of splitting* is denoted by *p*_*s*_, and may be calculated in a time interval by the equation 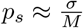where *σ* represents the total number of splits that occur in a time interval and *M* represents the total number of SEs that may or may not split through that time interval.

#### 3.1.4 Information transport in the SERD network - transport and storage

In the SERD network’s direct physical comparison, PPs represent the smallest element of matter in the system, therefore they are the constituents of particles and the foundation from which consciousness and observation emerges. PPs behave as fundamental microscopic observers and each have a unique view on what changes have occurred within the SERD network in which they exist within. They are the hubs upon which information is both updated and transferred.

In a given evolution (or branch) each PP has an *effective metric e*_*k*_(*i, j, t*) associated with it which defines the number of SEs between each of the pairs of PPs *i* and *j* that PP *k* is aware of at a given time *t*^19^. This is defined in more detail in [60]. This is the *observed state* of the system, and for a given PP, is determined entirely by the cumulative information that is incident on that PP over all time.

Each element action incurs a change to the topology of the network state at a given time. This topological change will affect the instantaneous *hidden* state of the system; however, it will not affect the *observed* state of the system for a specific PP. When an element action occurs it generates a bit of information that encodes the action on its bounding IGs. This bit of information is then passed through the system affecting its nearest neighbours through each TS. This results in a propagating and superposing wave front of information.

When considering the evolution of the information propagation of the SERD network with duplications, reductions and splits, there are two main types of information object to be considered; the *Propagating Information Packet* (PIP) and the *Propagating Structural Bifurcation* (PSB).

#### 3.1.5 Propagating Information Packets (PIPs)

PIPs are information storing objects which correspond to the propagating information of SE actions and exist in the IGs between elements. When an SE undergoes an action it creates a bit of information that is stored on the bounding IGs and then propagatesoutwards along the IE through time. Through each TS PIPs shift along one bounding IG of the SE to the other in their prospective directions of propagation. This process continues until the PIP reaches a PP where it both updates the metric of that PP but is also transferred to all other IGs connected to that PP. They superpose on one another via reductions^20^ and propagation is impeded by duplications. These are described in some more detail in [61, 60]. When PIPs reach a PP *i* but have not reached the central observing PP *k*, the information that has reached *i* from IEs that are not connected to *k* will be transferred accross the PP to all other connected IEs through one TS. **Therefore every TS a bit of topological information is emitted from a PP radially outward**. When fixing a PP at the centre this may be considered as radially inward. Fig. 5 shows how these bits of information superpose via the reduction of SEs as they propagate radially inward along an IE and how their propagation is impeded by the duplication of SEs.

**Figure 5.**
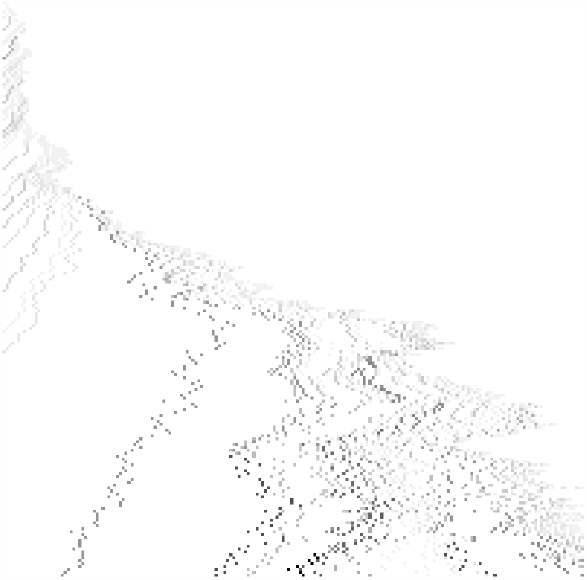
Space (horizontal)-time (vertical downwards) diagram demonstrating the production of PIPs along one IE of the SERD network. Information flows into the IE from the enclosing PP and then superposes with itself via reductions to produce PIPs.

It is these PIPs that in [61] it is argued could correspond to photons. They store information regarding changes to the intrinsic curvature of spacetime due to the IE around a PP that is not the COPP, and so when they inform a PP of the topologial change, by updating on that PP, consequently, the PP observes alterations to the spacetime topology. This observation leads to an effective change in the direction taken by PPs in the *time-iterated curvature minimizing*^21^ embedding of the *effective metric* described in [61, 60] leading to an effective force on the PP that the PIP came from^22 23^.

#### 3.1.6 Propagating Structural Bifurcation (PSB) propagation

PSBs are generated at PPs at the moment in time that the PP undergoes a *split*, as shown in Fig. 4. The number of PSBs generated is equal to the number of neighbouring IGs to the PP. They then propagate along the incident IEs at a rate of 1SE/TS, affecting each nearest neighbour each TS as displayed by Fig. 6, and continues until the PSB reaches a central PP as displayed by Fig. 7. A PSB is a propagating topological change in the *hidden* state of the network. That is, observing PPs are not *aware* of this structure until a PSB updates on them. A PSB is a wave front storing the information of a split, and sharing it with the neighbouring elements that it has not already propagated across. In this way it *unzips* the IE.

**Figure 6.**
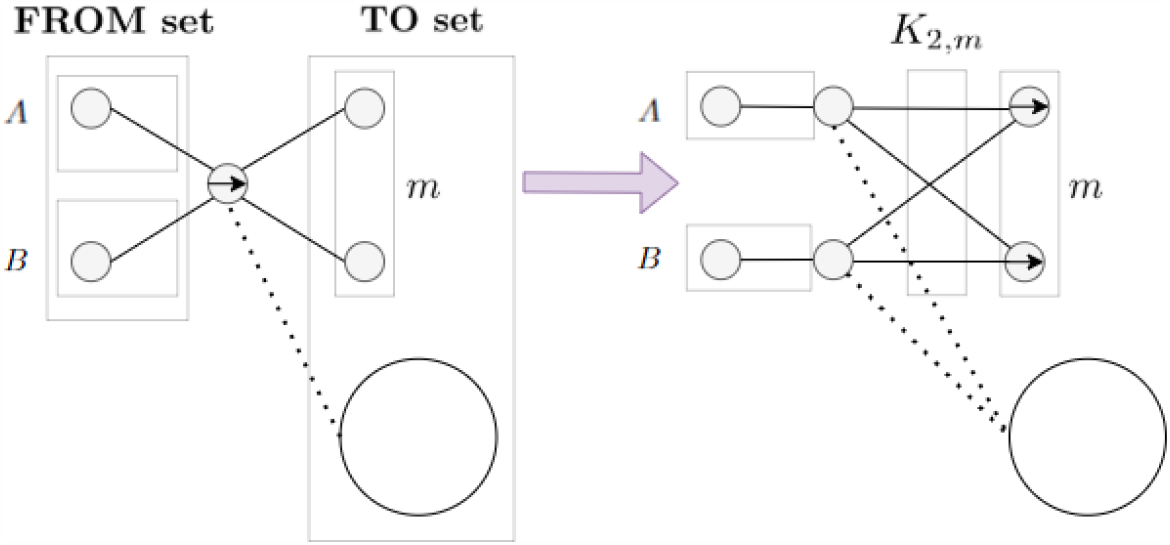
Diagram showing the update operation of *PSB propagation*, where *m* is the size of the edge set in the **TO set**. The PSB is centred on an IG called the host IG and has a **FROM set** associated with it. The **FROM set** defines all elements (SEs and PPs) that the PSB has already affected. The **FROM set** defines the **TO Set** from the incident element set of the host IG that the PSB is centred on. At each TS a new SE and IG pair are created for each element in the **TO set**, replicating any information stored therein. The PSB also replicates itself onto the bounding IG of each SE in the **TO set**. The **FROM set** may be partitioned into two non-overlapping subsets *A* and *B*, which may contain SEs or PPs. PSBs do not propagate accross PPs. The dotted lines represent adjacency.

**Figure 7.**
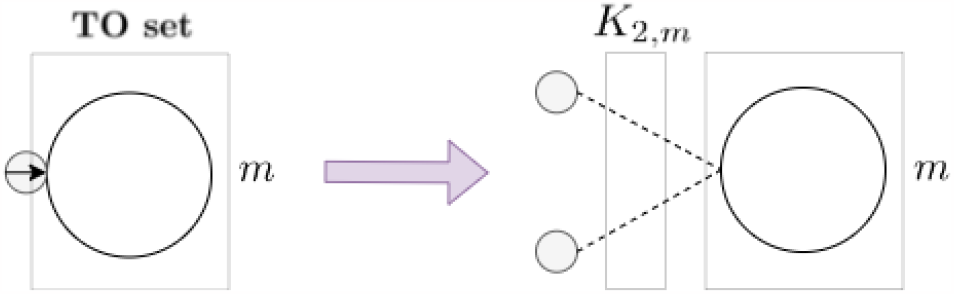
Diagram demonstrating the termination of a PSB on a single PP, or multiple PPs. This is the last step of a PSB, once the PSB reaches the PP the host IG splits in two and the PP becomes *aware* that a split has taken place.

PSBs are centred on an IG and have a **FROM set** and **TO set** associated with them. Since the IG has a neighbouring element set, only the information of the FROM set is required to be stored on the PSB object. They propagate along SEs and update on PPs. When the PSB propagates it generates a PSB for each SE that it propagates along, the PSB being centred on the next IG along. In this way PSBs may replicate themselves when they pass other PSBs propagating in the opposite direction as seen in Fig 8.

**Figure 8.**
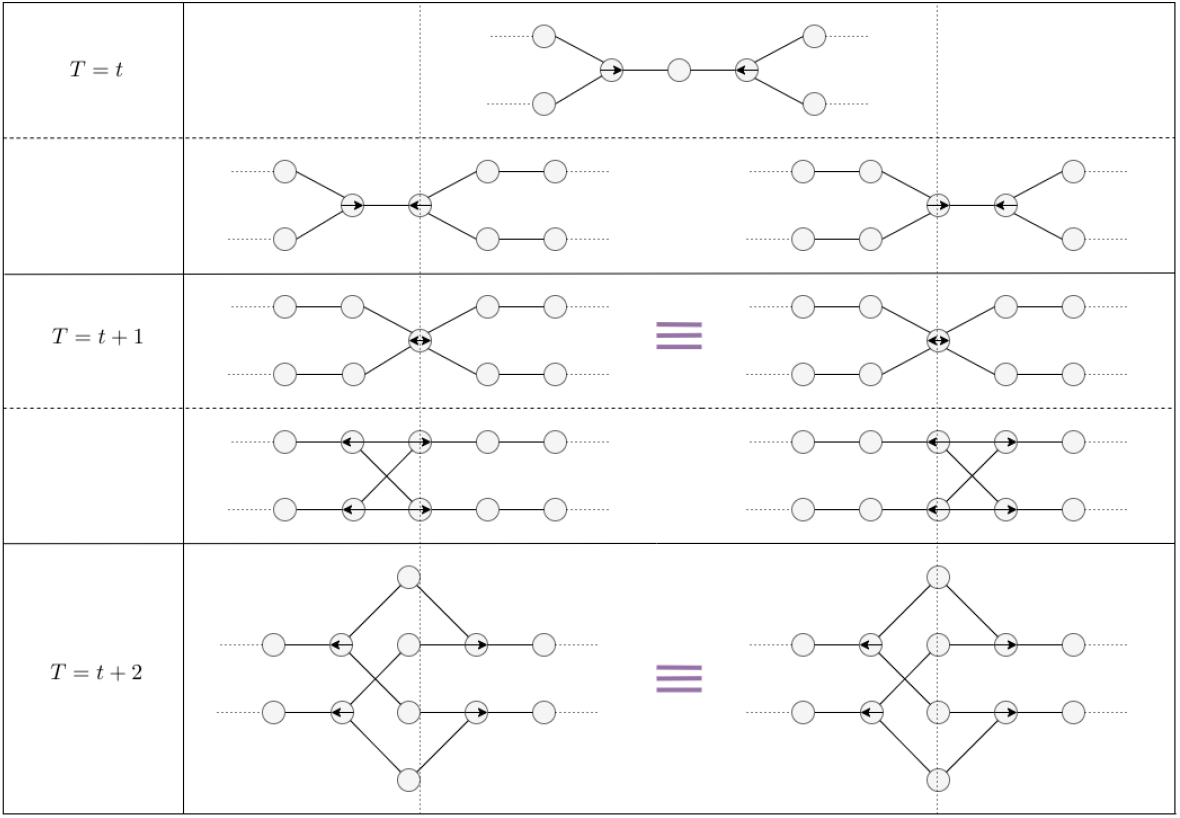
Diagram demonstrating the confluence of a simple PSB cross over. The implication here is that the output state is not affected by the order of update operations.

##### Definition 3.12.

*A* ***PSB set***, *B*(*t*) *is a set of all PSBs that exist within the SERD network at a given time t. This maybe an array of PSBs within the code or abstract as the information of the PSB is actually stored within the IG set*.

##### Definition 3.13.

*A* ***PSB index set***, *B*_*in*_(*t*) = (*n*_1_, …, *n* _|*B*(*t*) |_) *is a set of addresses that allows for access of the host IG of a given PSB. This set is run through linearly when PSB propagation is undergone*.

##### Definition 3.14.

*A* ***PSB*** *b* ∈ *B*(*t*) *is a mathematical object of the form b* = ((*P* 1, *S*1), (*P* 2, *S*2)) *where P* 1 *and P* 2 *are the PP subsets that the PSB will propagate away from and where S*1 *and S*2 *are the SE subsets that the PSB will propagate away from*.

Fig. 6 shows how a PSB at each TS resides on a host IG. At the end of each TS it then transfers the information of the PP split, in the form of topological structure of the network^24^, along the IE towards the COPP creating a spanning-tree radial topology represented clearly by implementations in the following sections^25^. Through each TS a PSB will sever the connections between two sets of elements in the **FROM set** *A* and *B*. From Def. 3.14 and Fig. 6 we have *S*1 ∪ *P* 1 = *A* and *S*2 ∪ *P* 2 = *B*.

#### 3.1.7 Note on the order of update operations

It is important to clarify that there is a separation in the order of non-deterministic *element actions* and deterministic *information propagation operations*. That is, in a TS, all *element actions* occur followed by *information propagation*, there is no overlap between the two types of operations. While element actions are nondeterministic and generate a *multiway system*, propagation operations are fully deterministic and occur simultaneously after all actions have occurred. Actually, PIP and PSB propagations occur instantaneously and simultaneously, both sharing information with their nearest neighbours. In computational time this is not necessarily the case since each information transfer must be performed in some order since PSB propagation changes the topology of the network^26^.

### 3.2 SERD network as a model for discrete space-time and information through the cosmos

It is important to make clear that the SERD network model is a new hypothetical model for a discrete space-time and is not established in the literature. Implementations and investigations into the behaviour of this model are still in thier early stages; however, there are a number of meaningful motivations and physical comparisons that provide some justification of the use of such a model^27^.

#### 3.2.1 Philosophical motivation

The SERD network model arises, as described in [61], out of a fundamental philosophical motivation. Having a strong philosophical motivation for rules provides intrinsic value to the model and is a question that is not addressed in other similar models for discrete space-time [42]. In this view there is a justification of the specifics rules, and structure, of the SERD network model.

#### 3.2.2 Emergence

The rules and fundamental elements of the SERD network emerge out of the philosophical motivation. Out of the rules and elements emerge complex structure. The endowment of elements with elementary awareness leads to information propagation, which then leads to the emergence of PIPs and PSBs. It will be shown later on in this paper that these rules generate biological topology, as well as inviting cosmological comparisons.

#### 3.2.3 Emergence of photon-like propagating information packets

The SERD network, via its dynamic information propagation mechanism has been shown to generate propagating information packets which behave in a similar manner to photons. They travel at an average finite speed of 1SE/TS, or 1 *plank length*/*plank time*. They update on matter particles instantaneously, creating a jump in the energy state of the particle^28^. It will later be shown in this paper that these photon-like information packets are redshifted under the expansion of space.

#### 3.2.4 Emergence of cosmological horizons

The information transfer of PIPs, and the effect of update on COPPs, leads to constraints of the expansion rate of the radial size of the observable universe [61]. That is, the metric that is defined on a COPP may not expand faster than the speed at which light, or information, can travel. This leads to separated *pocket universes* similar to those described in [90, 91, 92]. In *section 4.4* evidence will be presented to support the hypothesis that the combinded effect of PSB propagation and exponential inflation naturally leads to causally separated regions and information horizons, namely *pocket universes*. These are not only horizons relating to the observed radial size of the space, but the matter content aswell. We shall see how entire separated pocket universes may exist in the SERD network completely causally isolated from a chosen COPP.

### 3.3 SERD network as a model for mycelium growth and information transfer

The SERD network has a number of features that makes it suitable for exploring the mycelial morphology since, in addition to the cosmological and physical comparisons of such a model it also has many similar corresponding behaviours to mycelium.

#### 3.3.1 Apical branching

Fig 9 gives an example of how PP splitting leads to *apical branching* in the SERD network model. When a split occurs it generates a new *non-Radial IE* inbetween the two daughter PPs. PSBs are created in all directions including radial (towards the COPP). It’s the propagating branch structure of the PSB along a radial sub-graph of the SERD network, along with some expansion due to SE duplication, which naturally leads to the generation of a *spanning-tree topology*. Apical branching and spanning-tree topology is therefore generated by a ***growth-only model***^29^.

**Figure 9.**
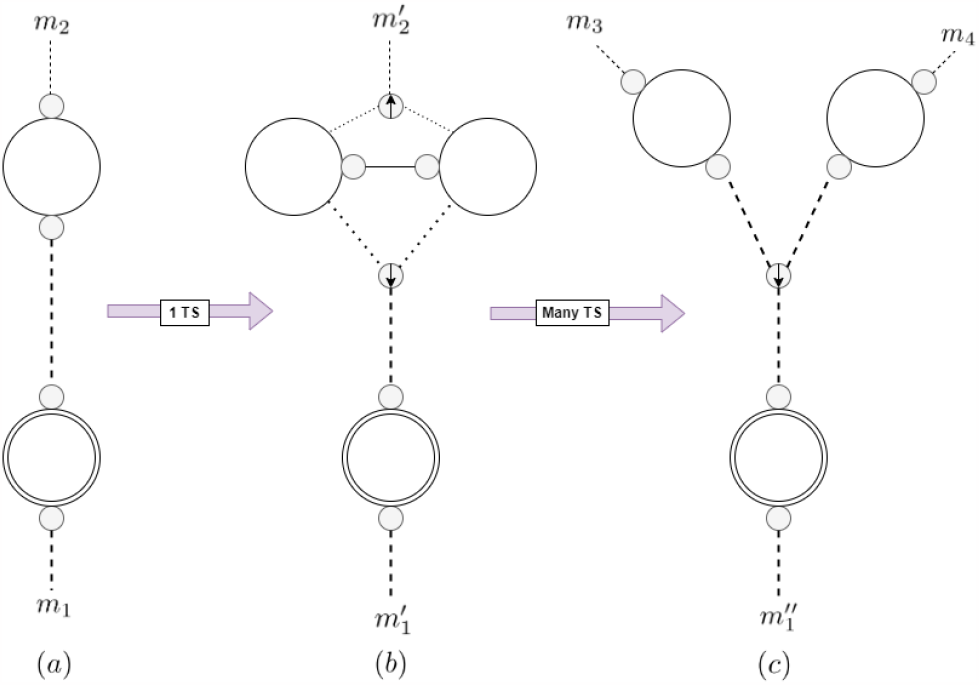
Diagram showing how splitting of PPs can lead to branching structure of the radial subgraph of the SERD network via PSB generation, with *m*_*i*_, 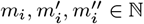 referring to the number of IEs connected to a PP excluding the connected radial IEs in the diagram.

#### 3.3.2 Sub-apical lateral branching

Fig 10 shows how an example correspondence to sub-apical branching mycelial morphology in the SERD network is hypothesised. There is an emergent negative pressure on the SERD network induced by the probability of an IE *fully reducing* that is significantly larger at smaller IE scales [61]. Since PSB propagation continuously adds a constant flow of 1SE per TS to the sub-IE branch it is possible for some proportion of sub-IE branches to be *reduced*^30^.

**Figure 10.**
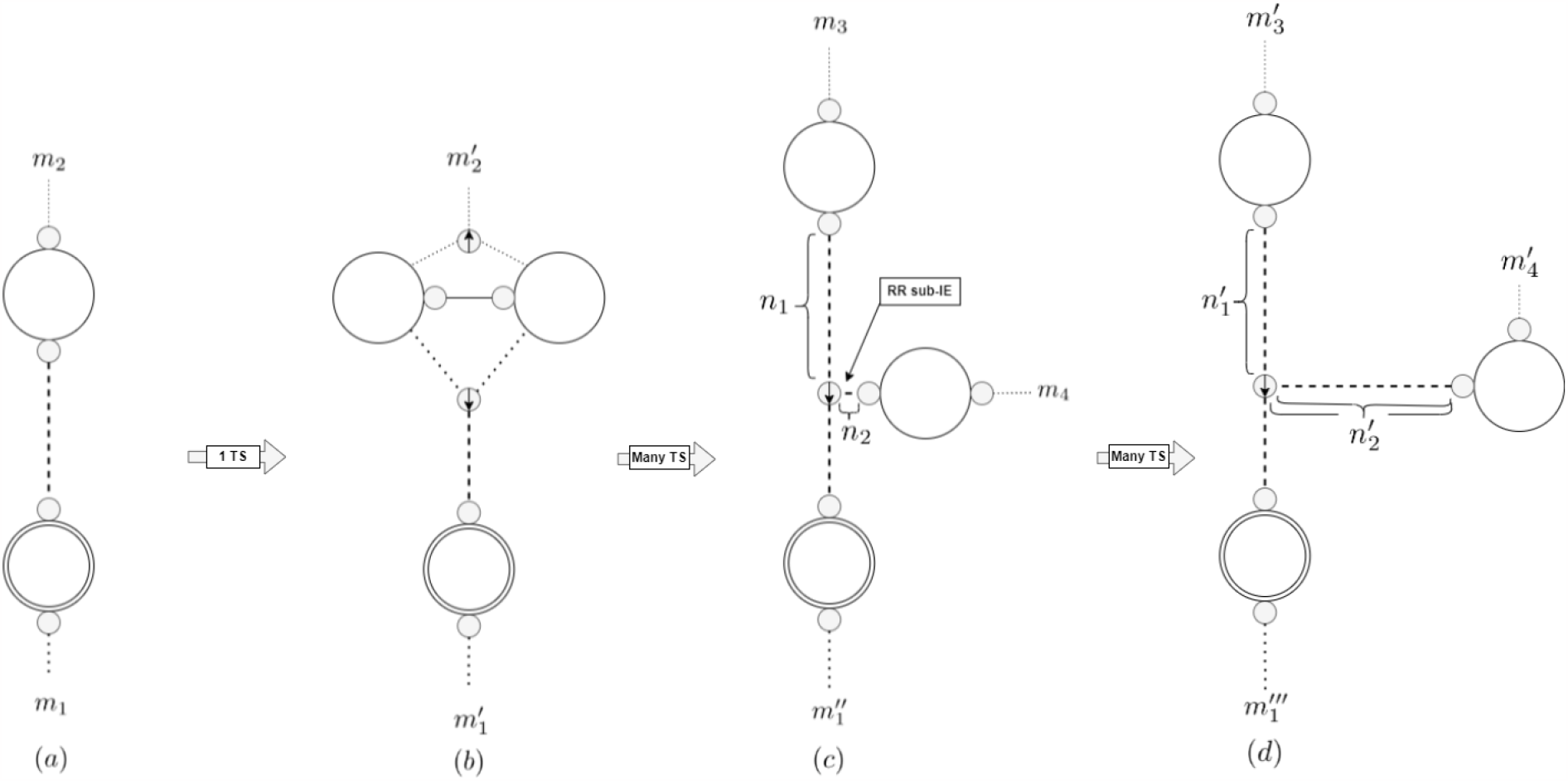
Diagram showing how subapical branching topology and morphology occurs in the SERD network model. With 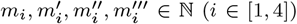 referring to the number of IEs connected to a PP excluding the connected radial IEs in the diagram and 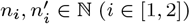 referring to the number of SEs between two IGs depicted in the diagram. RR sub-IE means *Reduced Radial sub-IE*. In this case 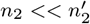

##### Definition 3.15.

A ***reduced sub-IE*** is an IE that bounds a PP and a PSB that is below a threshold size related to the increase in the probability of smaller IEs to fully reduce.

Once produced the *reduced sub-IEs* will propagate down the IE until they pass a *threshold scale* and the probability of full reduction reduces dramatically. At this point the reduced sub-IE is no longer reduced and acts as a sub-apical laterally branching hyphae.^31^.

#### 3.3.3 Anastomosis

In Fig 11 we see how *anastomosis* (hyphal fusion) occurs in the SERD network. A split occurs generating a reduced sub-IE. The PP that bounds the reduced sub-IE is connected to another PP via a NRIE. This NRIE then *fully reduces* causing the two PPs to become connected in an internal compactified PP space. This forms the point of fusion of the anastomosis. The condition, therefore, for anastomosis in the SERD network is that the fusion point must be between two PPs^32^, the NRIEs between which have fully reduced. This connects the two PPs, separated by a single IG, and therefor fuses the IE or hyphae together^33^.

**Figure 11.**
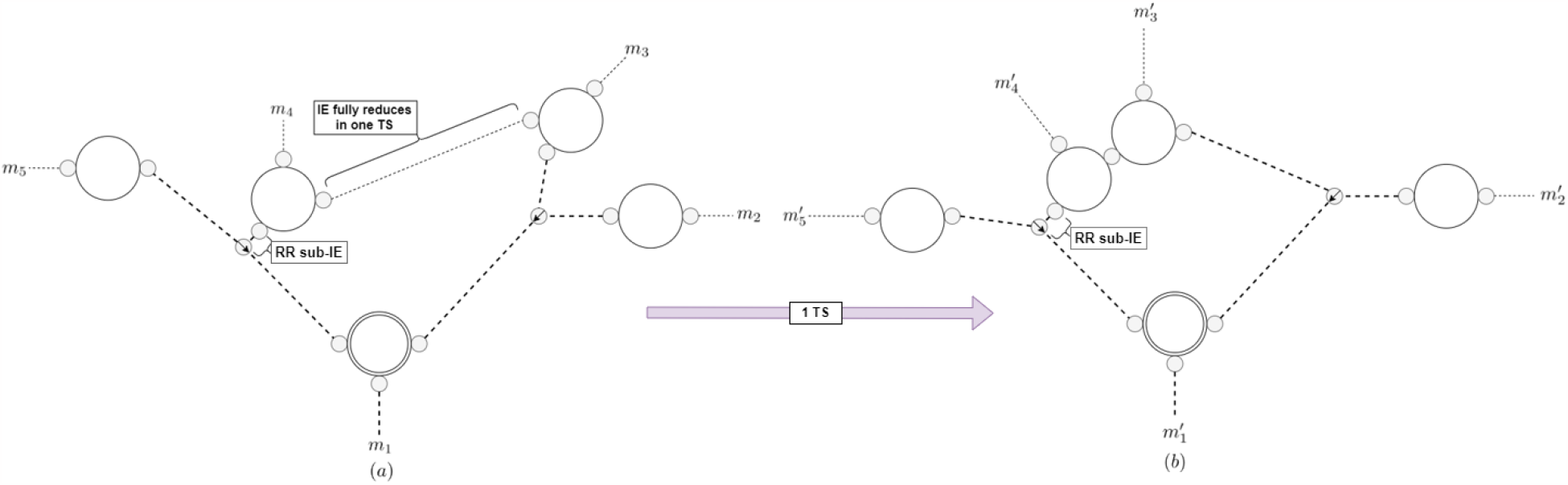
Diagram showing how anastomosis, hyphae fusion, occurs in the SERD network model. With 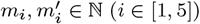 referring to the number of IEs connected to a PP excluding the connected radial IEs in the diagram. RR sub-IE means *Reduced Radial sub-IE*.

#### 3.3.4 Space elements duplicate like cells undergo mitosis or bacteria binary fission

One of the most fundamental behaviours of cells is that they replicate themselves under the process of mitosis when resources are sufficient and when a number of biochemical factors are activated. This replication, - or duplication -, process takes place everywhere in nature continuously. In the context of the SERD model’s mycelial comparison, and in comparison with other biological systems, SE duplication corresponds, in some very fundamental sense, to this behaviour. The growth of the SERD network is driven by the process of SE duplication, where an SE, replicates itself through a TS, becoming two from one. This process is highly reductive and does not involve the internal complexities incurred in a cells mitosis cycle. Nevertheless, it results in spanning branching tree morphology of a growing fractal network, and in the most general case follows a very similar modelling procedure, the growth rate being directly proportional to the quantity involved. In other words there is *exponential growth*, at least of the *hidden* state^34^. This is particularly the case when considering a *growth-only model* where we take *a*(*t*) ∼ *e*^*Ht*^, and how this process bears very striking similarities to the the exponential growth phase of cell or bacterial growth.

This is not entirely the case for filamentous fungi since when in the growth and foraging phase - a phase somewhat analagous to the initial inflation stage of the SERD network - unlike in the case of the SERD network, the rate of mitosis of filamentous fungi is not exponential, and the majority of cell replication occurs at the hyphal tips, rather than anywhere along the hypha, resulting in linear rather than exponential growth behaviour. The process of hyphal growth is more complex since it incorporates the generation of new hyphal wall from the Spitzenkörper vesicals at the tips and translocation of organelles and cytoplasm though intercellular pores, and does not follow the same cyclic mitosis process as in other organisms [93]. The hyphae feed the tips with nutrients needed and ready made organelles to increase the rate of apical cell replication [75]. In this sense sub-apical organelle generatation and translocation feeds the apical cell replication in a similar way to how SE duplication in the middle of an IE may influence the scale of the SERD network.

#### 3.3.5 SE Reduction is similar to cell death but with some key differences

Mycelium propagates through a substrate via cell mitosis, however this process may be hindered via a number of factors leading to cell death. This maybe due to resource scarcity, disease or grazers. In this process, the physical structure of the mycelium is interrupted and breakages in the hyphae are observed due either the cells bio-material being reabsorbed and translocated, disease, or consumption by grazers. In most cases this leads to breakages in the network topology. This is not the case in the SERD network. Although *reductions* result in the dissapearence of an SE, or cell, and result in decreasing the population in the system, SE reduction does not cause breakages in the IEs.

SEs are space itself, and so when reductions cause a change in the spatial separation between points, the neighbouring elements recombine to fill the gap^35^. This may happen in some cases in biological networks, however, biological networks are physical systems of matter which exist within the fundamental spacetime framework and are not the fundamental framework itself. When a cell dies, in the context of the SERD model it does not disconnect spacetime, it is in fact a translocation of matter and energy from one location in space to another. In this way reductions are somewhat analogous to cell death but not entirely isomorphic, and the distinction is that one network represents the a fundamental spacetime structure which is fully connected and cannot be broken and another network is a physical construct embedded within that space.

### 3.4 About the SERD network, mycelium and the cosmic web

In the introduction we cited [1] which made a claim regarding similarities in structure between slime mould *P. polycephalum* and matter in galaxies. This was extended to the cosmic web and evidence has been preposed of a link in matter formations at large scales in the cosmos and matter distributions in biological emergent networks [63, 64], in particular such as slime mould [6]. There is common ancestry and connection in morphology with mycelium [54, 62, 66, 67, 68, 94] and so similarities to slime mould may be found as well since we hypothesise the processes discussed in *sections 3.3.1,3.3.2* and *3.3.3*. In particular [64, 6] found that all matter, including dark matter, resides along long threads with void-like regions, similar to how biological networks form.

If *sub-apical* branching occurs in the SERD network then this implies the existence of reduced IEs, which in turn implies that long IEs tend to have PPs closely attached to them, like a chain, and so this may correspond to this matter formation on the large scales. Fig 12 shows how these chains of matter along an IE or hyphae may form^36^.

**Figure 12.**
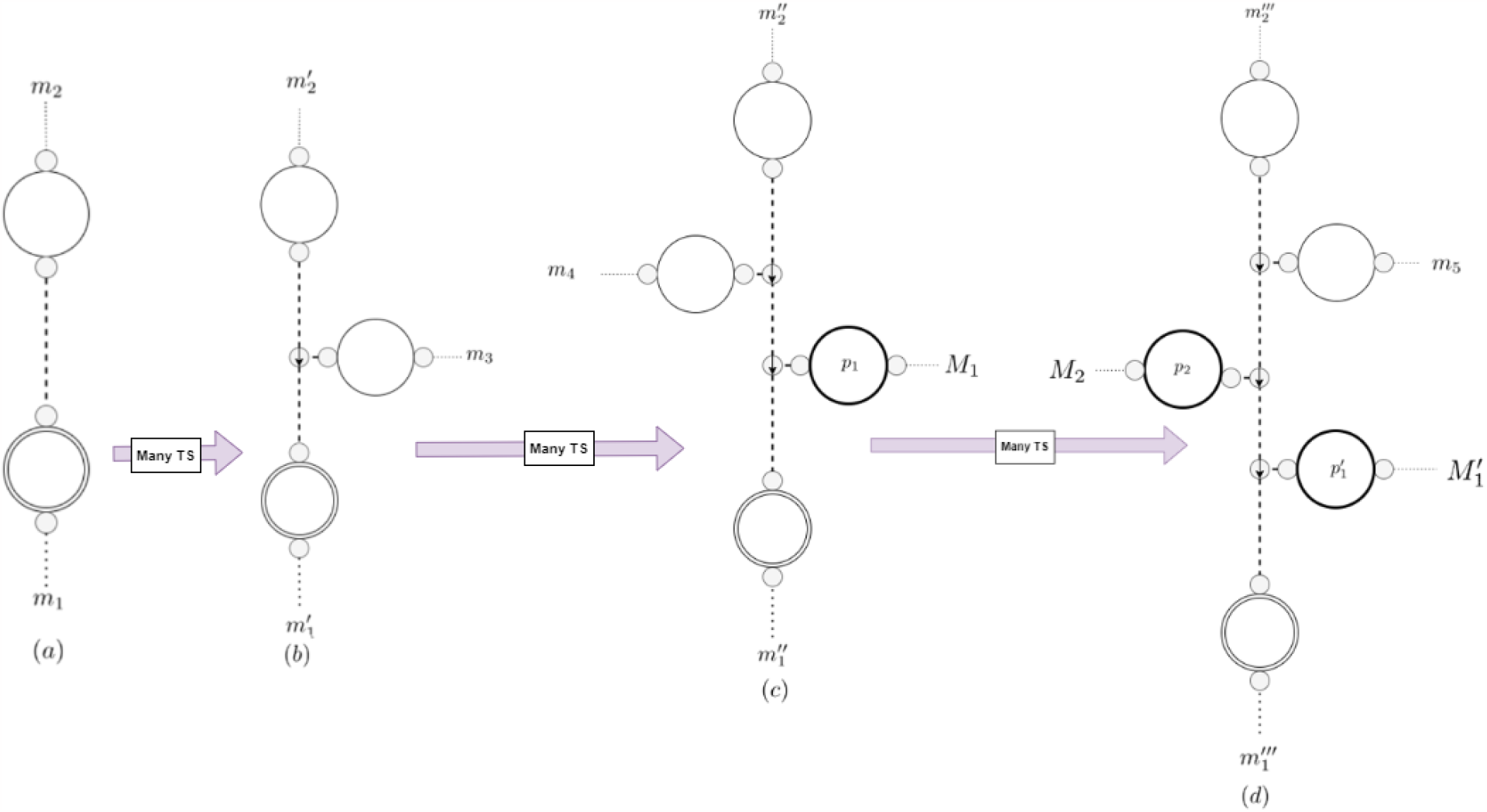
This diagram demonstrates how reduced sub-IEs between PPs and PSBs leads to matter being attracted to the IE or hyphae in the mycelial comparison.

### 3.5 The redshift scale-factor relation and direct computational testing implementation method

This section outlines the specific cosmological redshift scale factor relation that the SERD network will be tested against as well as the computational experimental architecture that leads us to the results provided in *section 4.4*.

#### 3.5.1 The redshift scale factor relation and the FLRW metric

The FLRW metric, derived from the cosmological principle of homogeneity and isotropism, has been successful at describing the evolution of the large scale structure and geometry of our observable universe. The FLRW metric is an integral part of the ΛCDM model. This model continues to corroborate observation and has recently passed a number of extremely high precision tests [95, 96, 97].

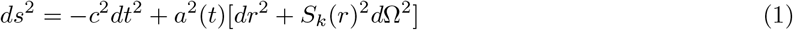

Equation 1 is of the FLRW metric, where *s* defines the space-time interval, c the speed of light and *a*(*t*) the scale factor. *dt, dr*, and *d*Ω represent polar spacetime coordinates. *S*_*k*_(*r*)^2^ is a parameter dependent on the curvature and radius which is defined as *r* for a flat spacetime. For the inward radial light-like trajectory of a photon this metric reduces down to *c*^2^*dt*^2^ = *a*^2^ (*t*)*dr*^2^, rearranging we get 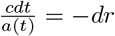 and then by integrating over the proper time and distance we end up with 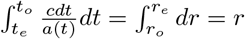. When the photon updates at the start and the end of one wavelength we have the same proper distance. 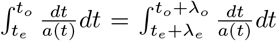. Then, adding the integral 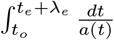 to both sides leads to 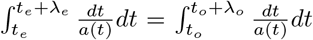. Then, using the fact that the update time for a photon is orders of magnitudes less than scales at which the scale factor changes we can bring out the scale factor from the integral and using 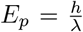, we get the following red-shift scale factor relation of the FLRW metric that will be used to test the feasability of the SERD network.

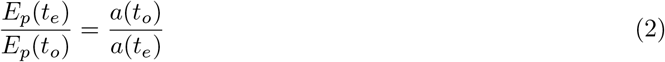

In 2, *λ, ν* and *E* represent the wavelength, frequency or energy of a photon respectively, at either the moment of emission *t*_*e*_ or observation *t*_*o*_. This relation tells us that the ratio of the energy of a photon at emission and observation is proportional to the ratio of the scale factor at emission and observation, and is one of the main cosmological comparisons of the SERD network model made in this paper.

#### 3.5.2 Energy information correspondence in the SERD network

In modelling this physical correspondence between information loss of a PIP and the energy loss of a photon, it is important to connect the information content of a PIP with the energy content of a photon. When a photon transfers energy to matter that energy is entirely absorbed by the matter particle, this may then drop by releasing another photon. Absorbtion of the energy of the photon increases the span of the wave function of the particle, causing a change to the particles momentum and the likelihood of it travelling further. A gain in energy is an increase in the distance a particle may travel in a given direction in a unit of time.

##### Definition 3.16.

A ***unit PIP*** is the array of numbers that is transferred to the IG of the radial IE neighbouring the *bounding PP*.

When reductions occur in the SERD network this causes information to superpose, concatinating the arrays of information, or unit PIPs, and resulting in an effective PIP magnitude.

##### Definition 3.17.

The ***effective magnitude of a PIP*** is the number of *unit PIPs* which constitute it.

A bounding PP releases a steady stream of unit PIPs, corresponding to information of SE actions from non-radial IEs, at the connected end of the IE at a rate of one unit PIP per TS. These unit PIPs then either spread out or superpose, via SE duplication or SE reduction respectively, as they propagate along the IE, see Fig. 5. Reduction-induced superpostions have the effect of bringing together a larger number of unit PIPs, increasing the *effective magnitude* of the PIP. This is much like enforcing a number of TSs to exist in the same TS on update, increasing the maximum distance able to be traversed and increasing the intrinsic energy of the particles state proportional to the magnitude^37^. **We therefore make the correspondence between the *PIP magnitude* and the *energy of a travelling photon***.

#### 3.5.3 Method to model observed expansion induced redshift in SERD network model

Cosmological redshift is observed as a decrease in the energy of propagating photons. In the SERD network PIPs propagate along IEs towards a COPP. We make the connection that the process by which PIPs update on the COPP and contribute information to the observed state, is the same - or similar - process as the process of observing photons^38^. Therefore, in this case, we are primarily focussed on the one dimensional single IE system.

Fig. 13 provides a space-time diagram of the experimental architecture of the implementated test of the redshift scale factor relation, with results presented in *section 4.5.2*. The initial input state of two IGs bounding a single initial SE is used for the first tier of a two-tier Monte-Carlo simulation^39^. Each tier consists of a number of *sample runs*^40^ and run for *t*1 and *t*2 TSs repsectively. The array stores a number *α*(*t, x*) ∈ N to represent the *effective PIP magnitude*^41^. We may think of *α*(*t, x*) ∈ℕ as representing the space components of the *effective PIP magnitude* array that exist on the IGs of the IE *α*(*t*) = (*α*(*t*, 0), *α*(*t*, 1), …, *α*(*t, x*), …)^42^. There are two *phases* to a TS in a run. First the *action phase* where SE actions are realised. ***Duplications insert a 0 into the array*** while ***reductions sum the magnitudes of two neighbouring IGs and then removes the other*** .^43^ Then information input and update occurs by removing the first array element, as updating on the COPP, and appending a 1 to the bounding end of the array to represent the transfer of one unit PIP from the bounding PP in that TS. This has the effect of information propagation along the array. A *dilated time* is calculated by keeping track of the total effective PIP magnitude that has updated on the COPP since the start of the run. It is this dilated time which alows us to extract the scale factor at time of emission *a*(*emission*) of a given observed PIP.

**Figure 13.**
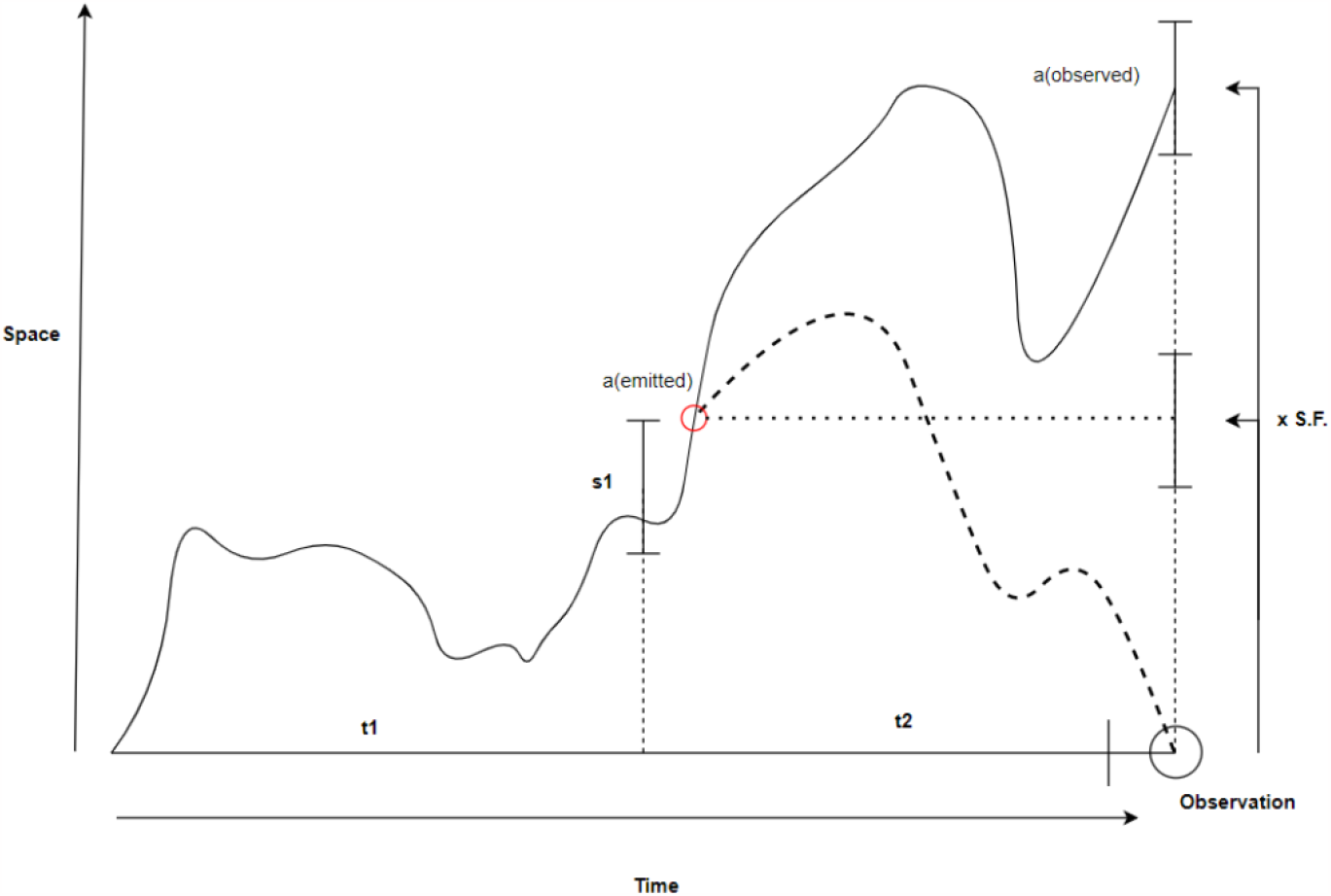
Diagram representing the structure of the experimental implementation to test for PIP-photon correspondence via the redshift scale-factor relation. The black line shows one possible evolution of the scale factor, the dotted line show the path of a PIP in the inflating system.

##### Definition 3.18.

The ***radial dilated time*** *t*′ ∈ ℕ is the observed time of a bounding PP which the COPP observes.

##### Theorem 3.1.

*The* ***radial dilated time*** *is equal to the sum of the number of unit PIPs that have updated on the COPP since the creation of the* 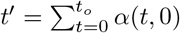, *where t*_*o*_ *is the global time at observation*.

*Proof*. All incident information on a bounding PP is transferred to the neighbouring IG in the IE connecting to the COPP in the form of a unit PIP every TS. This PIP encodes all information of the observed microstate of the bounding PP. The total PIP magnitude observed is equal to the total number of unit PIPs observed by the COPP and therefore equal to the number of TSs observed of the bounding PP from the point of view of the COPP. Thus, we may think of the total observed PIP magnitude, since the initial split and creation of the IE, as the ***radial dilated time*** *t*′ of the bounding PP.

In Fig. 5 it is possible to see the space-time evolution of a single IE, the input flow of *unit PIPs*, the denser PIPs that are generated via SE reductions and how these PIPs update on the COPP. A PIP, or photon, encodes topological information transferred from the bounding PP, or emitting particle, at the time of emission and therefor so long as we store an array of all scales at each time *a*(*t*) for each sample run and the dilated time *t*′ it is possible to retrieve the scale factor at the point of emission and at the point of observation. Fig. 13 shows how the Monte-Carlo simulation is split into two tiers:

##### Tier 1

The *scaling tier* evolves for *t*1 TSs to grow the scale of the system from the singe SE state. Only states with outputs within a given range at a scale of *s*1 will be stored in the input set. When the scale is within the range at TS *t* = *t*1 the dilated time *t*′, the scale factor history *a*(*t*), and PIP array *α*(*t*1) are stored for inputs for the second testing tier^44^. This generates naturally scaled input states.

##### Tier 2

The *scaled tier* evolves for *t*2 TSs and tests redshift scale factor relation at a given scale. A number of sample runs are implemented using the input sets from the last tier as randomly chosen inputs values. Two subsets of outputs are stored from this. When the run is within a given range from the final TS the PIPs at the COPP contribute to the extracted variables. At the point of update on the COPP the dilated time *t*′ and evolution history *a*(*t*) is used to extract out the scale factor at time of emission of the PIP *a*(*t*′) = *a*(*t*_*e*_). The *PIP magnitudes* that were updated get totalled and the total sample size counted and stored for each possible value of *a*(*t*_*e*_). This is implemented in, both the static system, where the system does not grow, and for the growing system where the scale factor is increased by some scale factor ratio. From this data an average PIP magnitude is extracted for specific scale factors at emission for both the static IE *Ē*_*p*_ and the growing system *Ē*_*p*_*′* within a given range of the final TS.

It is important to state that the input scale *a*(*t* = *t*1) is not the same as the scale at the point of emission of a PIP. The scale of the observation at the end of the tier 2 sample runs is chosen based on the scale at the point of emission of the specific PIP, not the enforced scaling that gives the input for tier 2. This method was used to implement the results presented in *section 4.4.2*.

## 4. Results

In this section we present results implemented from computational implementation and experiment, as well as mathematical analysis of, and computational tests for the SERD network’s cosmological comparisons. The SERD network is multifaceted and maybe implemented in various different levels of completeness^45^. Developing a complete SERD network model implementation with all included features is, and has been, an incremental process and has been throughout our research into the network. This involves first implementing a network with splits, duplications and reductions, but no PSBs or information propagation [59], then implementing information propagation but with no splits or PSBs, [60], and then further refining the model and providing a more in depth look into the radial growth characteristics of the model, emergent from the dynamic information propagation [61]. Each paper has built on the previous one. In much the same way, the results in this paper provide new insights into the biological and c osmological comparisons of the model. A fully workable code that implements SE reductions alongside PP splitting and PSB propagation had not been successfully developed yet. This means that we have a good model for exploring the larger-scale effect of growth on the information content of PIPs and testing it against cosmological models and some aspects of the mycelial comparison. However, it also means that specifically *non-apical lateral branching*, and *anastamosis* are not observed^46^. Nevertheless, the *spanning-tree structures* and *non-trivial connections* are observed.

This section is split into four subsections. In *subsection 4.1* we introduce the growth only model. In *Subsection 4.2* we show some graphical implementations of the *growth-only model* with *spring electrical embedding* and discuss correspondences with mycelium networks^47^. In *subsection 4.3*, building on the results from [61], we show evidence of Hubble horizons, resulting from the exponential spatial expansion and information propagation of the SERD network. We give a definition of the *Hubble parameter* in the context of the SERD network model and show graphical evidence that *branch clusters* of the *spanning-tree* topolgy are causally separated pocket universes. In *subsection 4.4* we present results that support the the hypothesis that PIPs are in fact photons in the phyical comparison, as the energy/information loss due to space expansion matches that of a photon as it propagates through an expanding universe. This not only supports the PIP photon hypothesis, but it also provides some direct physical comparison between the SERD network expansion and cosmological observation in the form of the FLRW metric and the corresponding *redshift scale factor relation*.

### 4.1 Evolution of the growth-only model and interacting PSBs

As stated at the start of this section, the results obtained when implementing PP splits, PSB propagation and SE duplication is the called the ***growth-only model***. In this model reductions do not occur, instead the system evolves monotonically and exponentially on average^48^. This model is an incomplete model of the SERD network since it does not include reductions; however, it works well for exploring the effect of expansion on the dispersion of propagating information, visualizing causally separated regions of the SERD network. It also demonstrates some correspondence to the fractal spanning-tree structure of mycelium and connection of sporulation locations^49^. We will see in *section 4.3.3* that interaction of PSBs generates non-trivial connectections between PPs.

Fig. 14 shows one branch evolution of this model. In this model the first few evolutions create the blueprint of what the final state will look like; then, once the space between PPs has grown to a specific threshold, PPs are constrained to a finite number of localized *PP clusters*^50^. When reductions are included this effect will not occur since IEs will have the capacity to reduce in size, resulting in a more dynamical evolution, when compared to the growth-only model.

**Figure 14.**
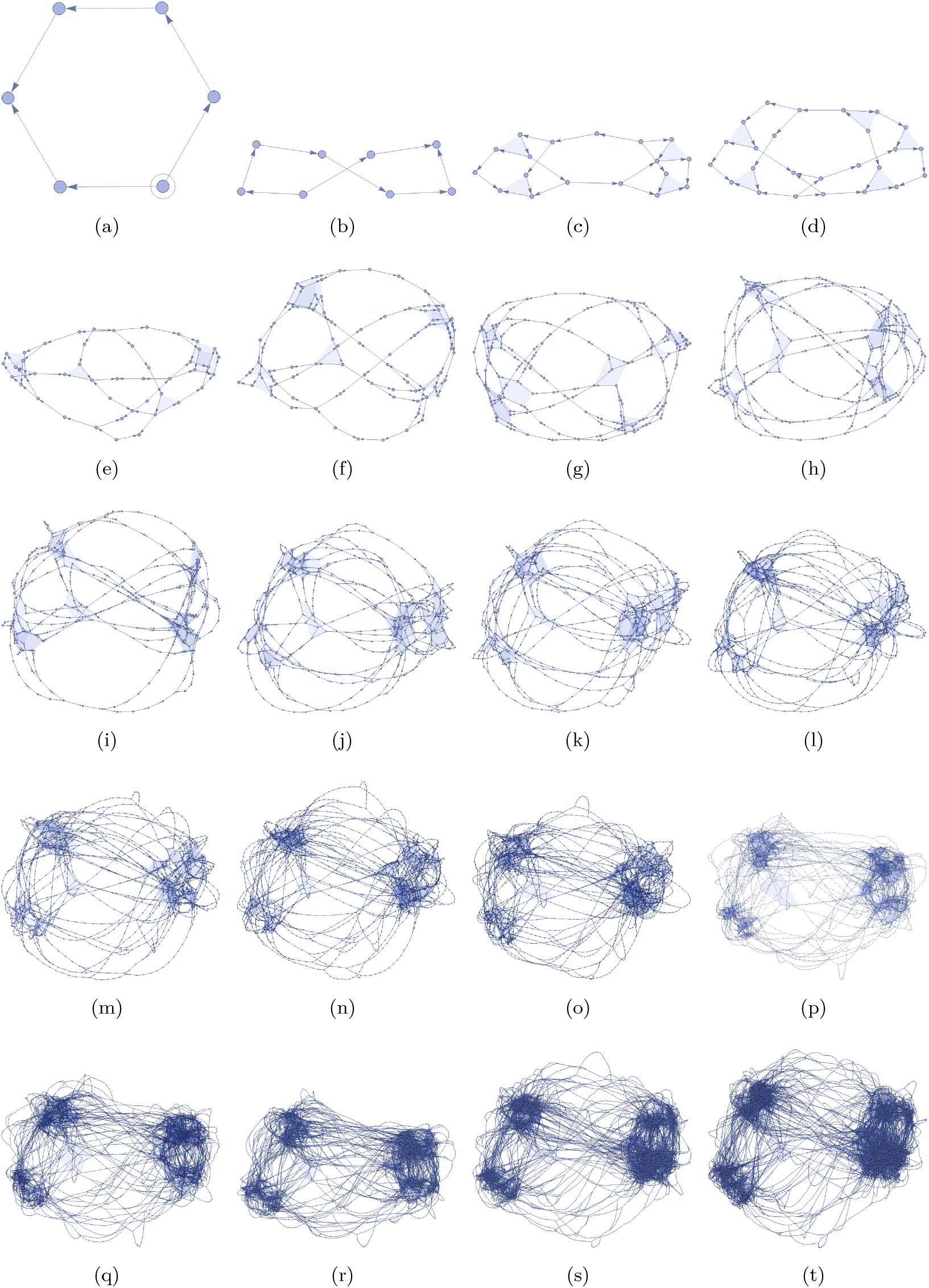
Snapshots of one branch of the growing SERD multiway system with time intervals of 1TS, with (a) at *t* = 1 from the initial single SE, two PP, state. The PPs are represented by the blue hyperedges and it is clear to see why these PPs may be referred to as hubs as the IEs converge on them. This network evolves via the *growth-only model*. The images provided show spatial expansion via SE *duplication* and creation of PPs via PP *spitting*. A Wolfram Model spring electrical embedding procedure is used. These figures are implemented using a *slow-growth model*, where 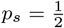 and 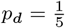.

**Figure 15.**
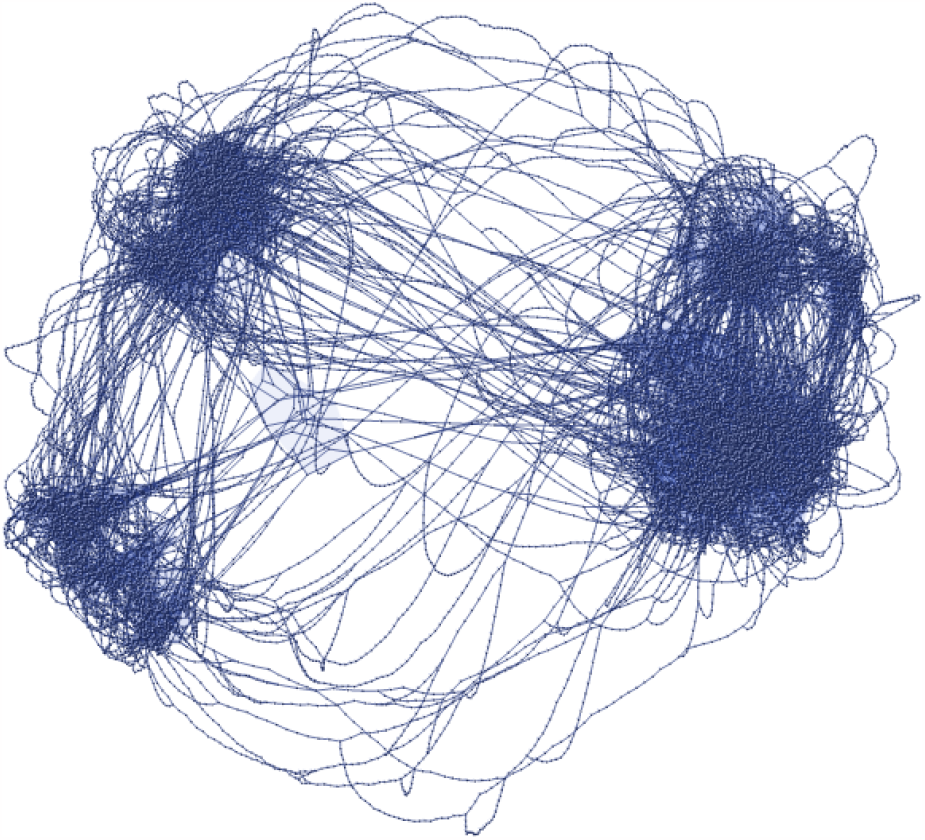
Global state of the SERD network implementation after 20TS of the SERD network evolving with SE *duplication*, PP *splitting* and PSB *propagation* embedded in space with *spring electrical embedding*. This SERD network state was generated using a *slow-growth model*, where 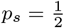 and 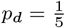.

Fig 16 shows another example of an embedding of the network at *TS* = 20. This is the parent state of the multispore radial subgraphs shown in Fig. 25, 26 and 27. Fig.17 shows a state where duplication is slowed, as can be seen the PPs form PP clusters or clouds much like those described in [59], where we propose that the number of PPs in a cluster is proportional to its mass.

**Figure 16.**
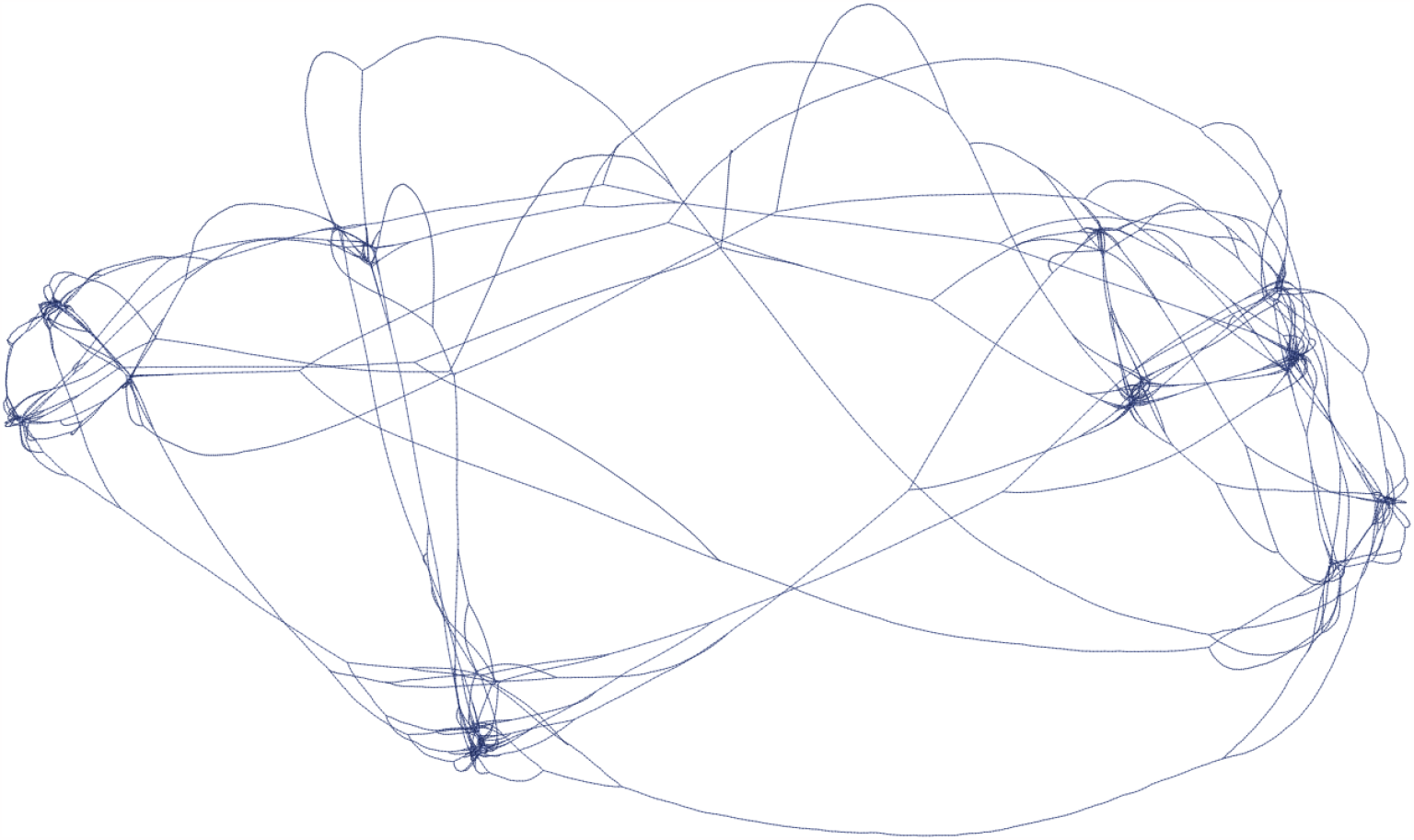
Global state of the SERD network implementation at *t* = 20*T S*, with 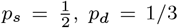 and *p*_*r*_ = 0. System embedded with *spring electrical embedding*.

**Figure 17.**
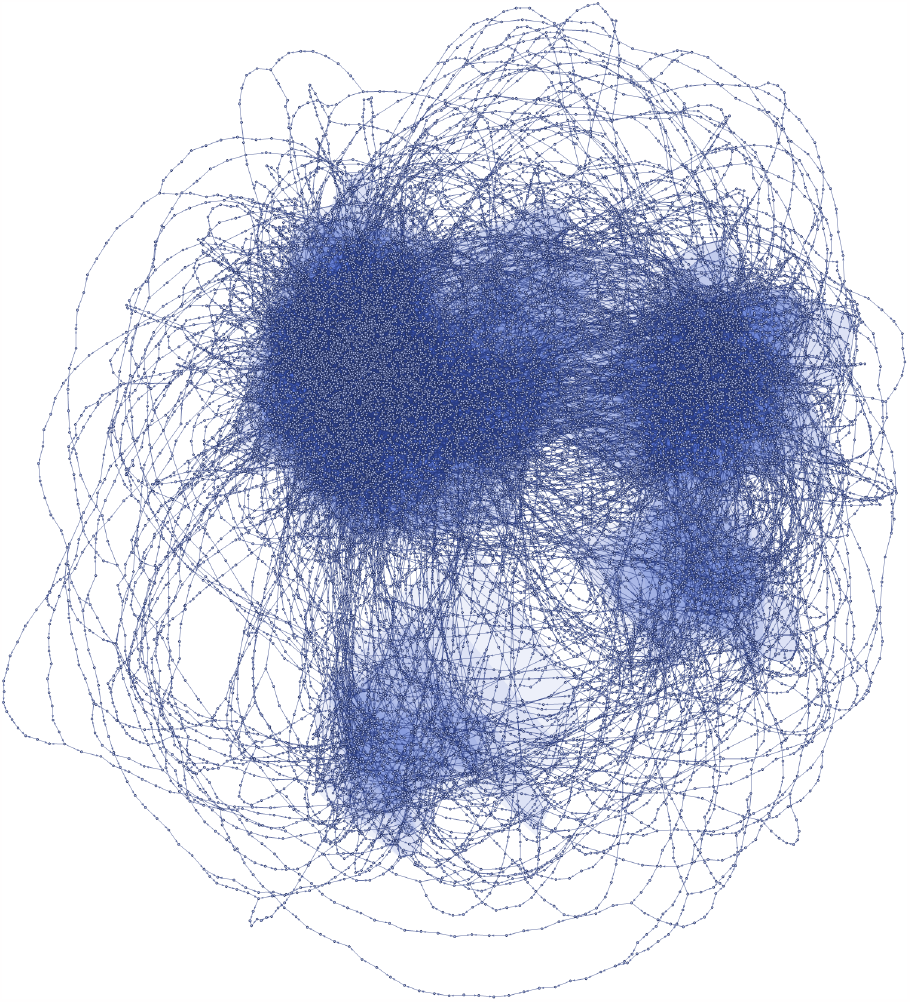
Global state of the SERD network implementation at *t* = 30*T S*, with 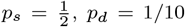 and *p*_*r*_ = 0. System embedded with *spring electrical embedding*.

**Figure 18.**
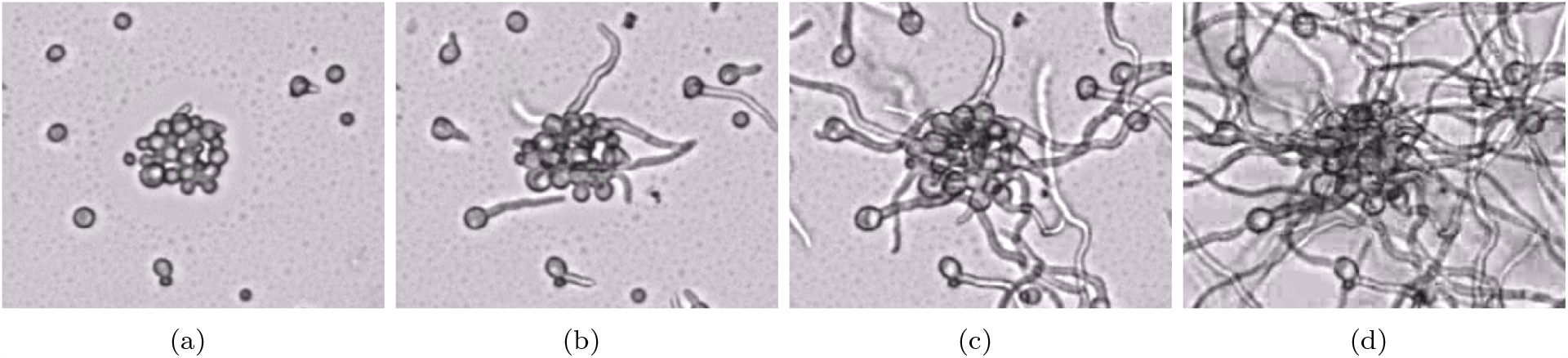
Clusters of spores germinating in a sample with other spores each frame taken every 4.5 hours. Published with kind permission of Professor Han A. B. Wösten, Wieke R. Teertstra and Ella M. Schunselaar, University of Utrecht

**Figure 19.**
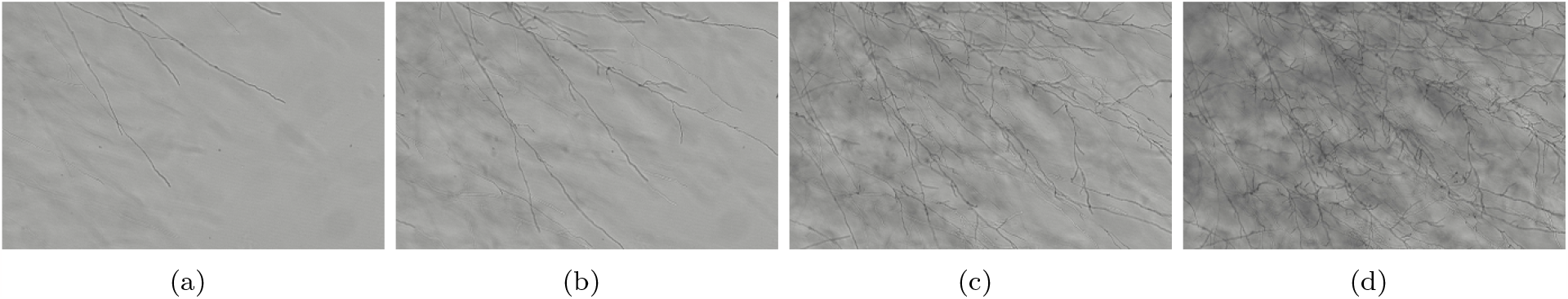
Clips showing the evolution of mycelium growth. The linear propagating behaviour can be seen as well as lateral branching. Each clip was taken at increments of 3 hours. Published with kind permission of Professor Han A. B. Wösten, Wieke R. Teertstra and Ella M. Schunselaar, University of Utrecht

### 4.2 Structural similarities in emergent topology of SERD network and mycelium

This section will discuss the mycelial comparison of the results implemented. It is very important to understand that - due to implementation of the *growth-only model*, as opposed to implementation with SE reductions - a number of direct mycelial comparisons described in section 3.3 will not be realised in this paper, namely *sub-apical lateral branching* and *anastomosis*. These behaviours will be observed once reductions are also included in the base code of the SERD network.

#### 4.2.1 The germination phase

The SERD network starts with a single PP in much the same way that mycelium begins with a single spore. This PP splits and generates a single IE between two PPs, treating one PP as the COPP and the other as a bounding PP we can see the correspondence between the initial single IE state and the germinating phase of the mycelium, where a single germ tube^51^ protrudes and then elongates. This can be seen in the first few TSs of Fig. 22. In that observed simulation the second PP immediately branches to form a PP and *sub-IE*, which in the comparison would correspond to apical branching in the neighbourhood to the spore which is not the case in observation of germinating fungi spores. There are two reasons for the difference between the results. First, there is the possibility that the branch taken leads to an arbitrary large growth of the first 1D IE, this may be unlikely as we increase the scale in space and time, however it is still a possible outcome and maybe considered as a valid input state. This result would correspond more to the comparison at the germination phase and any apical branching would occur further from the central spore location than PPs representing hyphal tips. Secondly since reductions don’t occur we don’t observe *reduced sub-IEs* and therefore no lateral branching is observed. When including reductions the multiple hyphae that would protrude from the spore location would correspond to *sub-IEs* that are either, connected to - or in the neighbourhood of - the COPP that break through the *reduced sub-IE threshold scale* introduced in *section 3.3.2*.

### 4.3 The foraging/growth phase

Figs. 20,21,24,22 and 23 all show the evolution of the radial subgraph of the SERD network. Dispite the lack of lateral branching and anastomosis due to the absence of SE reductions, a *propagating spanning-tree topology* is clearly visible in each instance.

**Figure 20.**
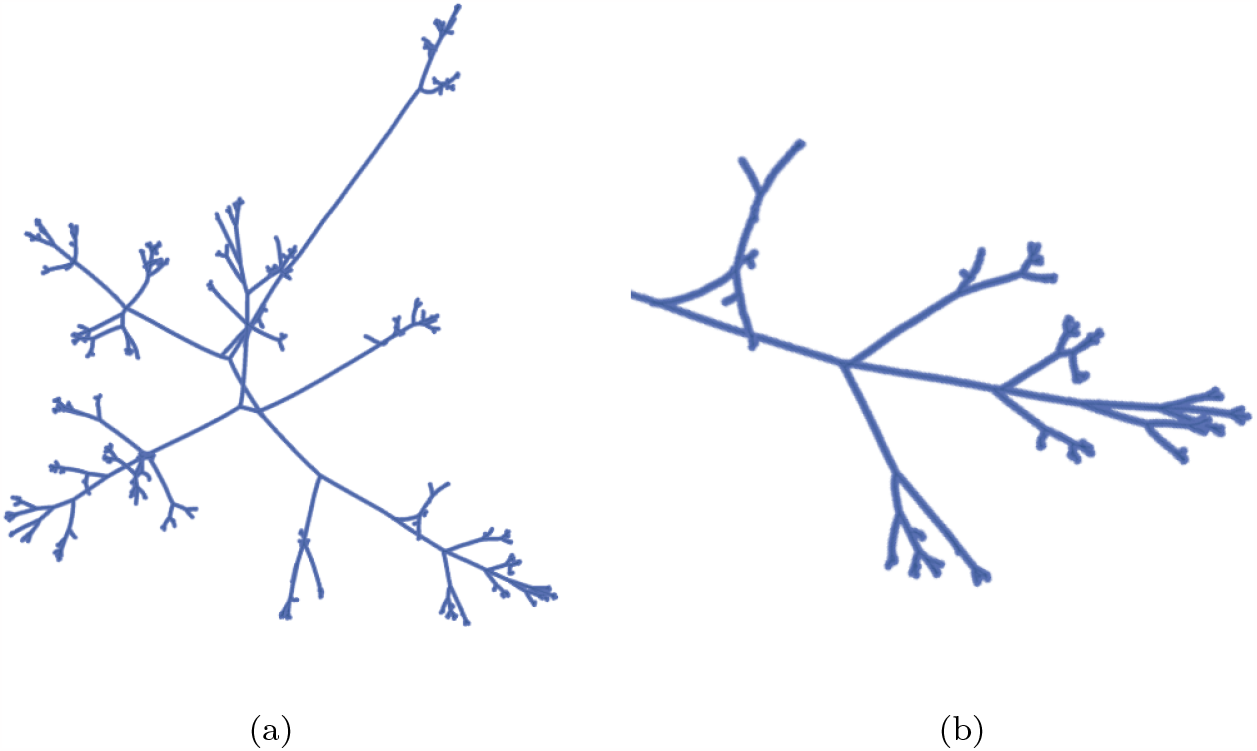
Evolved SERD network after 160TS of *slow-growth* under *spring electrical embedding*. At this scale it becomes difficult to distinguish branch points of the fractal structure and the occurrence of branch points of degree greater than 3 is clearly possible under SE reduction.

**Figure 21.**
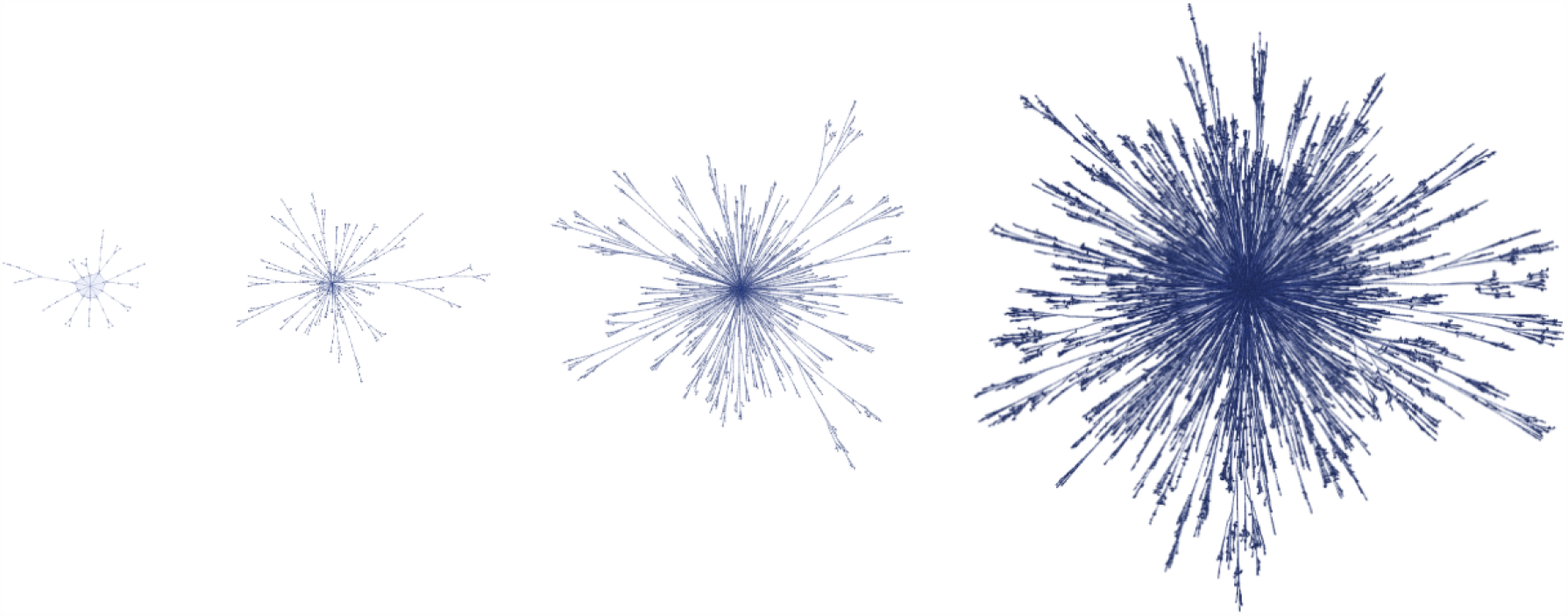
Evolution of a radial tree representing the growing radial subgraph of the SERD network. This evolution was generated with a sped up radial-only algorithm. Duplications, splits and PSBs were included, however reductions were not. These clips were taken at intervals of 25 TSs.

**Figure 22.**
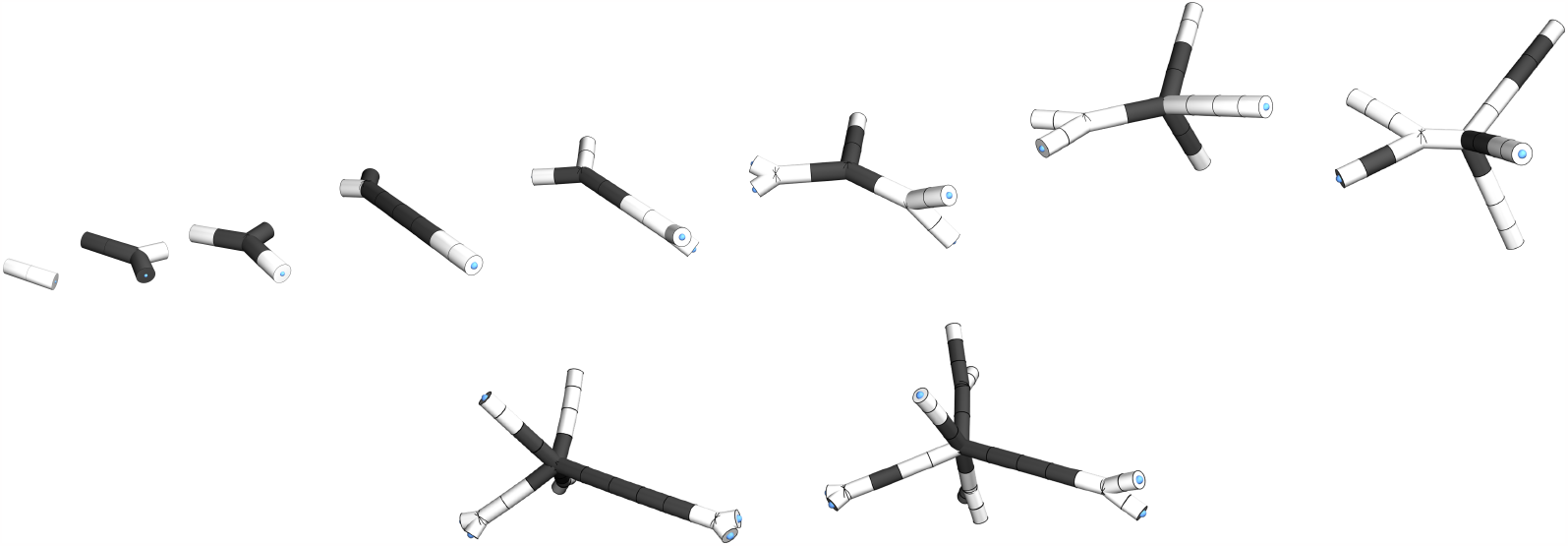
Evolution of radial subgraph of SERD network embedded with *spring electrical embedding* in the first 10 TS with inward information propagation. This graph evolves via the growth-only model. White cells represent an SE that is next to an IG that is storing a PIP.

**Figure 23.**
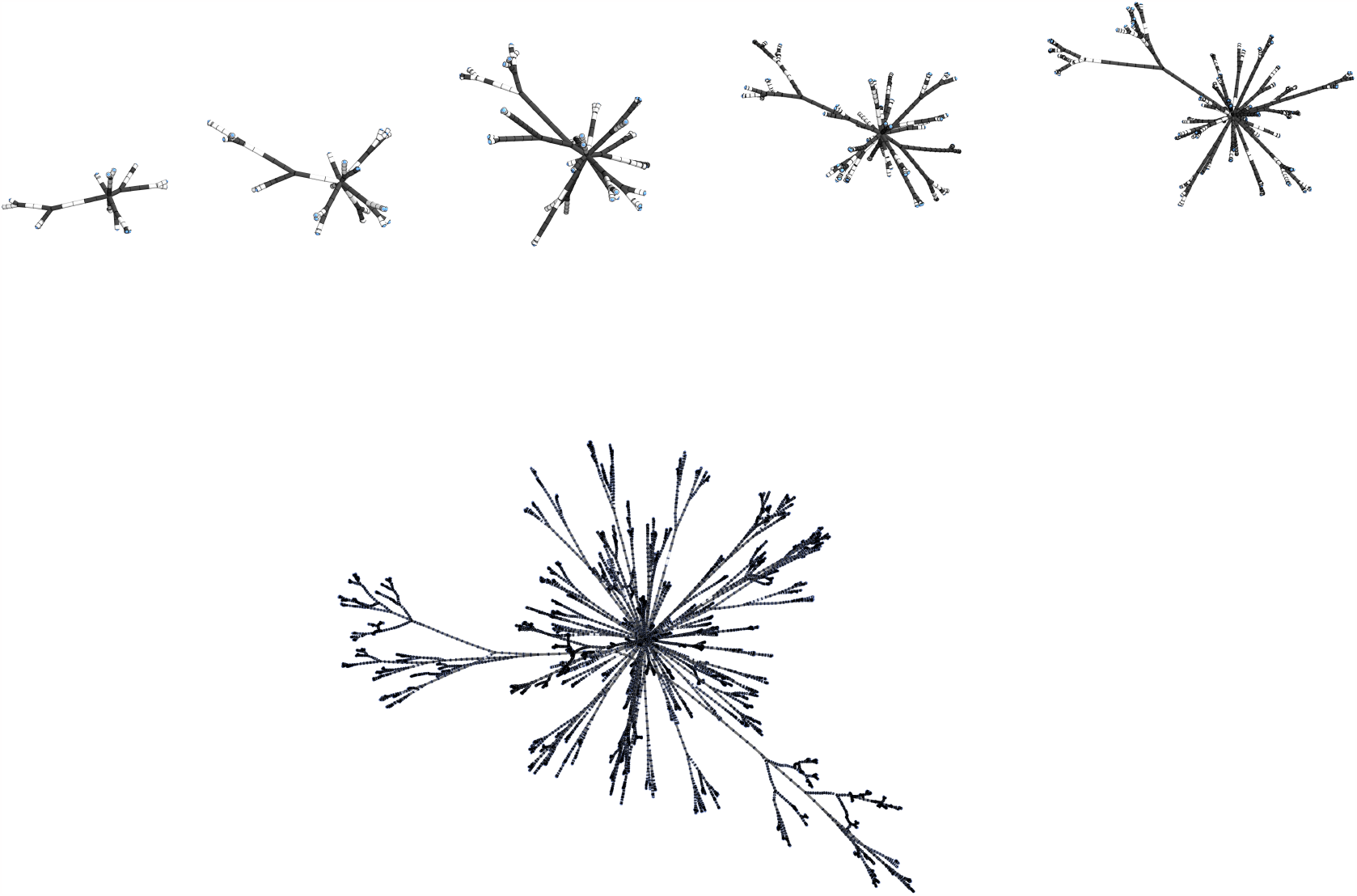
Evolution of radial subgraph of SERD network incorporating splits and duplications but no reductions. Embedded with *spring electrical embedding* from *t* = 12 to *t* = 20 with gaps of 2TS with inward information propagation. White cells represent an SE that neighbours an IG that is storing a PIP. The final graph is at *t* = 30.

**Figure 24.**
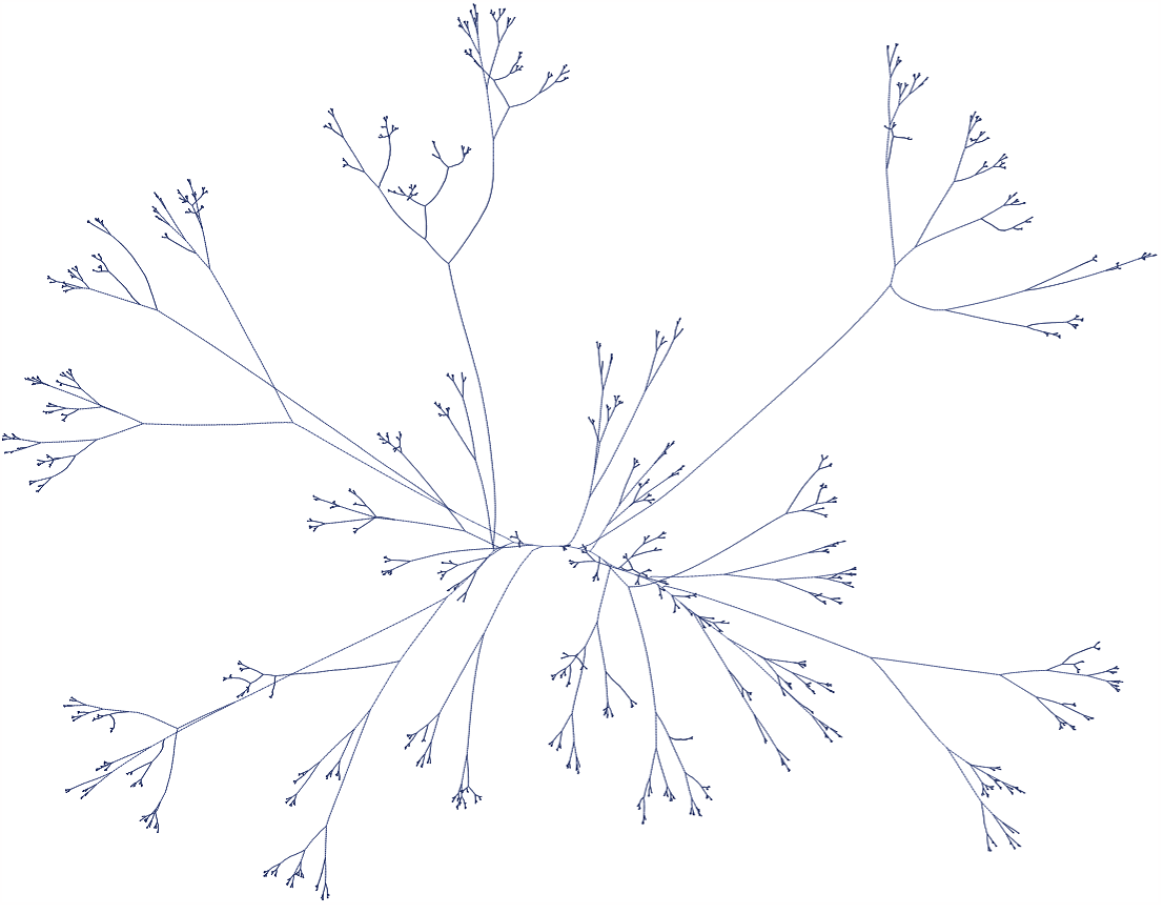
PP centred radial subgraph SERD network structure after 30 TS with *p*_*s*_ = 1*/*2 and *p*_*d*_ = 1*/*5. Split induced inward propagating structural bifurcation generates spanning-tree topology. SE reductions and nonradial IE are not included and so in evolution lateral branching and anastomosis would not be observed.

**Figure 25.**
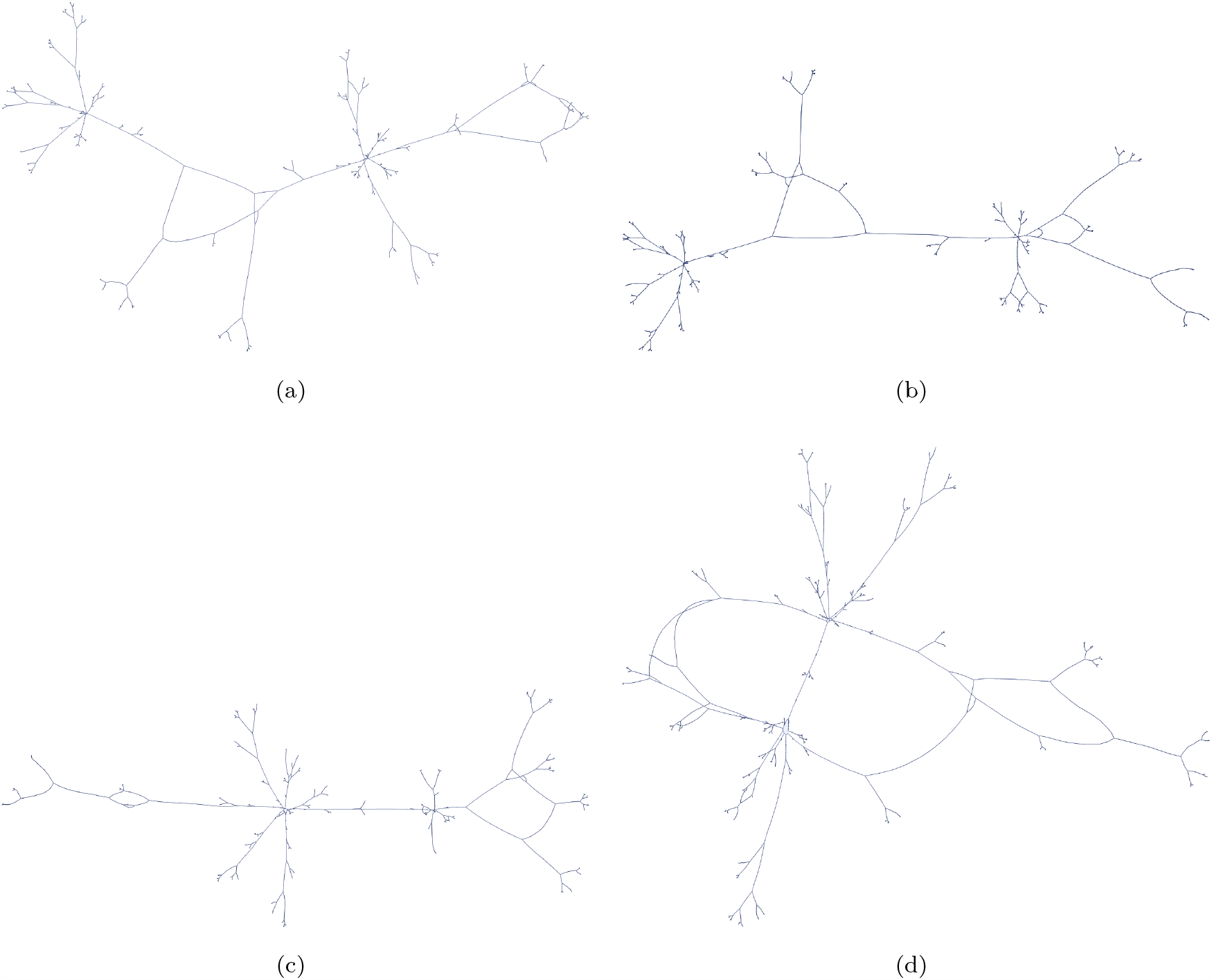
(a) to (d) show network diagrams of the connected radial subnetwork spanned by two PPs chosen at the 20th TS of the network state represented by Fig 16. The PPs can be observed as the two hubs and all ends of the IEs or hyphae. Nontrivial connections (more than one path for information to follow) between PPs is observed.

**Figure 26.**
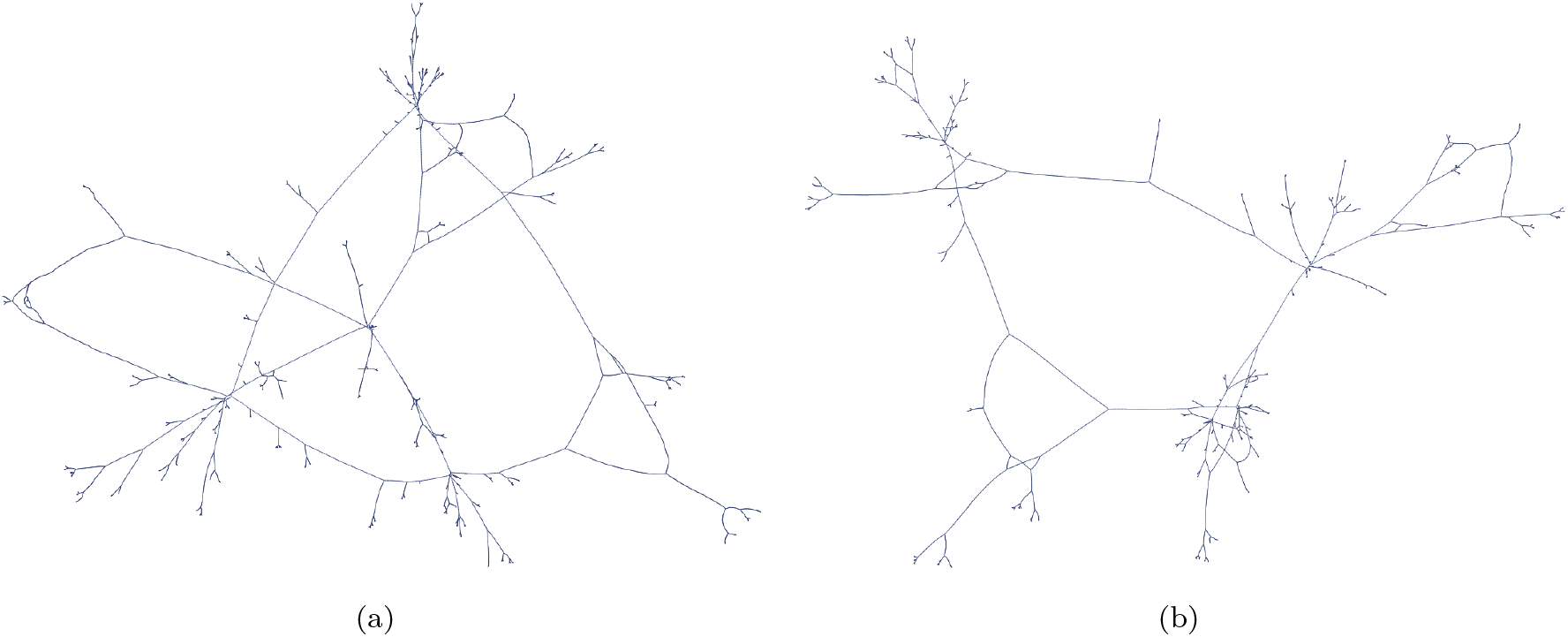
Radial subgraph spanned from four PPs (sporulation locations), taken as a choice of PPs from parent network displayed in Fig. 16. Interactons between PSBs can be observed as generating dynamic cycles. Information propagates in all directions along the network and so cyclic topology will result in self interaction of PIPs.

**Figure 27.**
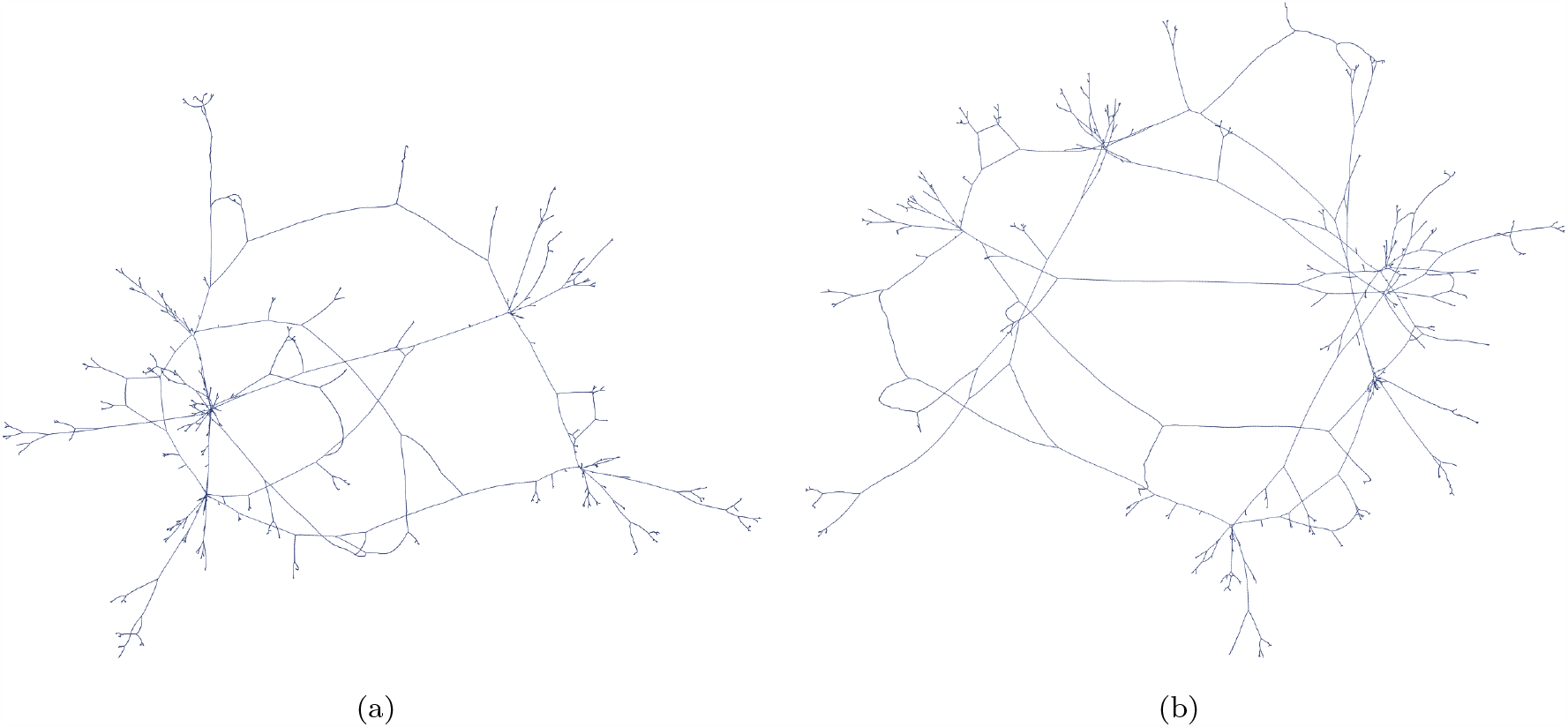
(a) and (b) show network diagrams of the connected radial subnetwork spanned by eight PPs. The PPs can be observed as the two hubs and all ends of the IEs or hyphae. Nontrivial connections are observed.

In each of these cases the COPP is constrained to not split to avoid outward propagating PSBs. This generates ineraction cycles with the inward PSBs. Exact spanning-tree topology is observed when the COPP does not split and we only observe inward PSBs^52^.

The PP’s splits generate apical branches, which propagate along the IE or hyphae. As the SEs duplicate this has the effect of impeding the progression of the PSBs propagation, which causes the branches to appear stationary - despite propagating at a rate of 1 SE per TS, as is case with mycelium. Therefore there is correspondece between the rate of production of new hyphal tips and the probability of splitting *p*_*s*_. The radial tree spans out in all directions as the network evolves, generating fractal mycelium-like topology.

While this network grows exponentially, mycelium growth tends to be linear in time and constrained by environmental factors. As demonstrated in [61], this exponential expansion is constrained within the observable horizon to be below the speed of light and so while the global unobserved system grows exponentially, the observed size of the system is constrained to grow linearly, with the horizon receding away at 1 SE per TS, or the speed of light. Therefore, there is both exponential and linear growth of the hidden and observed states respectively. It is also possible to ahmend how time is implemented in this network update model, affecting the growth rate and making a more realistic mycelial morphology.

Figures 19 and 20 shows some similarities. The lack of lateral branching in the *growth-only model* means that the majority of the branching tends to occur at the tips in the SERD model and since the probability of splitting is a global parameter^53^ we observe both fractal patterns and a shortening of the relative distance between branch points^54^. The fractal nature of the branching produces a greater fractal dimension around the tips of the IE, resulting in a greater number of IE further from the centre.

Figure 21 shows how when the network is allowed to grow to a large size and is embedded in 2 dimensions the it evolves in a similar way to foragin mycelium networks imaged extensively in [75]. The evolution in Fig. 21 shows the growing spanning-tree topology and the 360-degree spanning growth. However, as shown in [75], anastomosis and lateral branches are not observed and so updates to the base code including reductions will be required for further mycelial comparsions in this regard.

#### 4.3.1 Information transfer

Figs. 22 and 23 show not only how the SERD network evolves in the *growth-only model*, but also demonstrate, how information propagates along the IE from the bounding PPs inwards towards the COPP. This corresponds to how signals are passed from the apex through the network. In the global SERD network this information propagates in all directions along IEs, in a similar manner to how information is passed along hyphae.

At this stage this is still an early model and when reductions are included there will be both superposition of PIPs aswell as PIPs diverging and reconvening at branch points. Due to the expansion of the network from SE duplications the highest density of inward-propagating information is at the apexes, since that is the source of the information. It is then dispersed as the system grows. This is demonstrated in 1D with Figs. 32 and 33.

#### 4.3.2 The fruiting/sporulation phase

In the observation of radial subgraphs 23 and 24 a concentration of branching detail is observed at the apexes. In an qualitative observation, these small branch clusters at the end of the slender hyphae have an increase in complexity and fractal dimension. Fruits are produced at the ends of branches and apexes of hyphae in mycelium and so it is reasonable to see a correpondece to detailed apical branch clusters. At the end of each IE is a PP. We can consider that the apex PP is much like a spore. Fruiting bodies occur in the apical region of the mycelium and produce spores. In a similar manner PPs are at the apex of the IE, yet they themselves may also be the centre of their own spanning-tree radial sub-network, in much the same way that a spore may produce its own spanning-tree network once germinated.

The sporulation phase in the SERD network may be considered in the following manner; a mature SERD network has a large number of bounding PPs, the SE along one sub-IE connecting a bounding PP grows without it splitting, forcing the bounding PP to be some distance in space away from the COPP. The PP then begins to split generating its own radial spanning-tree network. In this way PPs have the capacity to act as apices of hyphae, hyphal hubs for a mycelium networks and the spore itself^55^.

#### 4.3.3 Multiple sporing locations

The SERD network is a fully connected network. It contains no vertices of degree 1 and no disconnected subraphs. Each PP generates its own spanning tree. These spanning trees connect and interact via the overlapping of PSBs creating non-trivial connections and cycles between PPs^56^. Figs. 25,26 and 27 all use Fig. 16 as their parent network. PPs in the SERD network always have at least 1 IE between them and so the PPs that are hubs in the subgraphs in this sections will never be disconnected from another PP.

Fig. 25 shows this PSB interaction between two PPs, while Figs. 26 and 27 show the radial subgraphs centred around two PPs. While this does not follow precise mycelial structure^57^ there are a number of interesting novel features of this emergent topology.

Firstly, the connections are non-trivial, meaning that if we were to consider information propagation in this network self-interaction of information may occur^58^.

Secondly, information propagation on graphs of this nature will give rise to unconventional computational processes and information processing.

The PSBs form cycles that span out until they reach a bounding PP at which point they update. This operation demonstrated by Fig. 7 clones an IG, the node, and so in this embedding it is perceived as a separation of the two sub IEs and so maybe perceived as two separate hyphae. For anastomosis to occur, when reductions are implemented in the SERD network model, bounding PPs must be included in the edge set of the radial sub-graph so that IE, or hyphae, can fuse with bounding PPs as hubs. If bounding PPs are included in the edge set these cycles will remain, even when the PSB updates on the bounding PP.

#### 4.3.4 SE reductions included radially

The inclusion of SE reductions in the SERD network would generate a more complete model. It would allow for both PSB and PIP superposition on IGs, leading to branch degrees of greater than 3. There would also be fully reducing IEs leading to compactified PP spaces and anastomosis similar behaviours. Figure 28 gives some preliminary view of implementations of these radial sub-networks^59^.

**Figure 28.**
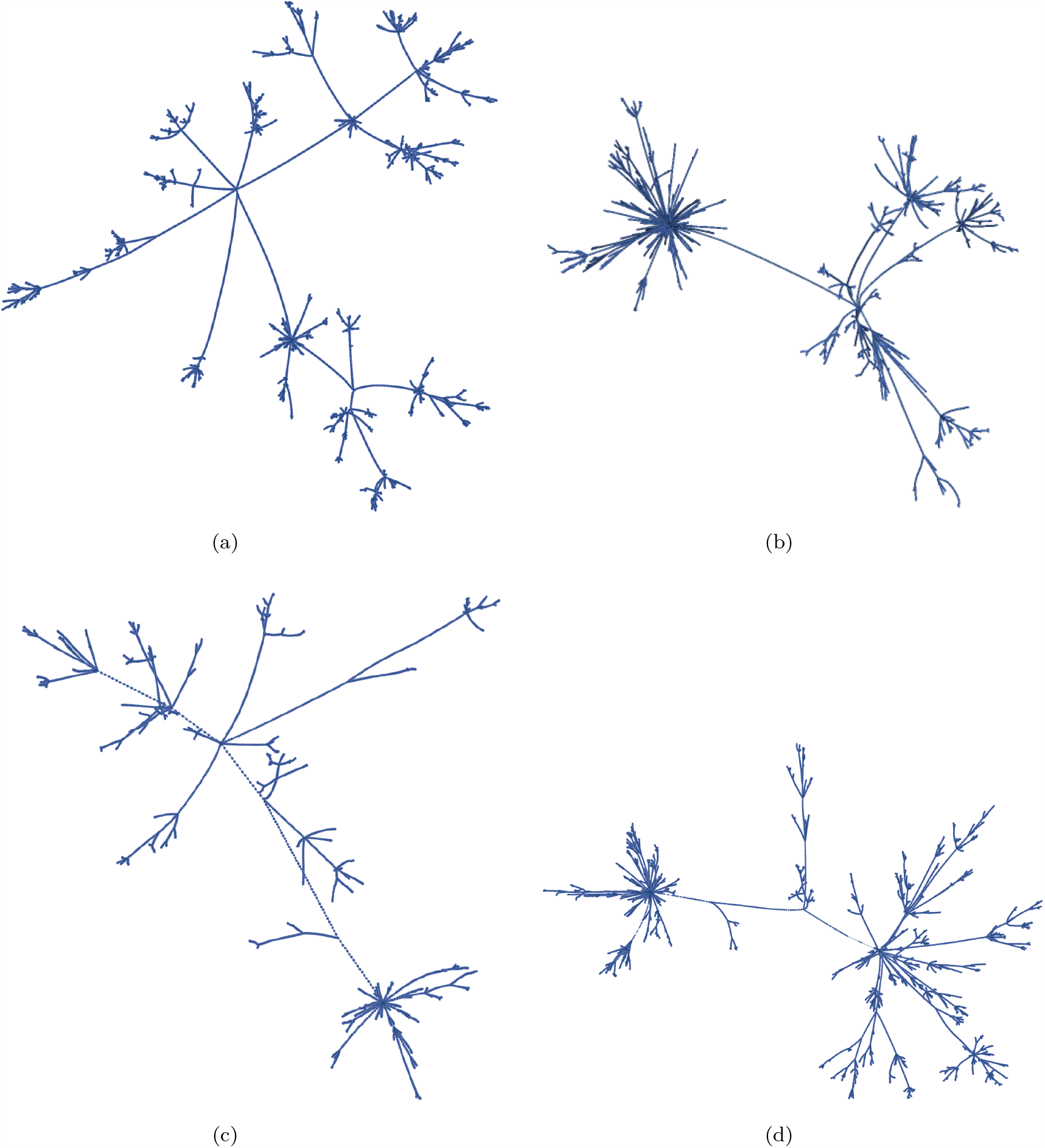
(a) to (d) show some example states of the radial subgraphs spanned from a single PP, with all hyphae ending on PPs. These states were obtained after 50TS with *H* = 1.06. Reductions have been included and so when two PSB meet they superpose generating PSBs on an IG nodes with a degree greater than 3. These were embedded with *elastic-spring embedding*. The evolution of these network structures with reductions has led to evolutions of smoothly embedded radial graphs. This will be investigated in detail in future papers, for now these images are just to give some idea of the possible radial topologies of the SERD network, and its relationships to biological networks.

### 4.4 Hubble horizons and branch clusters as pocket universes in an inflating space

In [61] evidence of emergent light-like limits on the observed radial size of the SERD network is presented. It is shown that the scale of the observed size of the system is constrained to grow linearly and proportionately to the propagation time that information has had to travel through the system and update on PPs. The limit on the observed scale was given to be the age of the universe and the distance to the observable horizon is the distance that light would travel in the universe’s lifetime. In these measurements only PIP propagation from SE actions were considered. In this section we will consider the causal horizons that affect the propagation of information from splits in the form of PSBs.

A PSB is very similar to a PIP in the sense that it propagates at a rate of one SE per TS, it may also *superpose* with other PSBs via SE reduction. What we show in this section is that the SERD network naturally generates these horizons, not just for PIPs but for PSBs also; that the size of the radius of these horizons is precisely 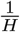 and then, given the results from [61] that 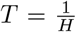, where *T* is a rough estimate for the age of the universe.

Firstly let us define what we mean by the Hubble Parameter in the context of the SERD network model. In cosmology the Hubble parameter is given by the ratio of the rate of change of the scale factor with the scale factor itself 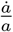, which ma ybe considered as either varying or as constant.

#### Theorem 4.1.

*The* ***SERD network Hubble parameter***, *through a given TS t, H*(*t*), *can be calculated as H*(*t*) = *p*_*d*_(*t*) − *p*_*r*_(*t*), *where p*_*d*_(*t*) *and p*_*r*_(*t*) *are the effective probability of duplication and the effective propability of reduction at a given time t, respectively*.

*Proof*. The rate of increase in the scale factor is the difference between the number of duplications and reductions 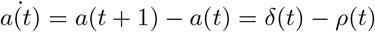. Therefore 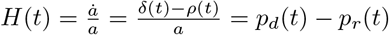. □

Using computational results from implementing a 1D growth-only model on the discrete space of SEs show in Fig. 29 and mapping light-like trajectories we extract a definition of the *Hubble radius* 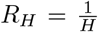, demostrated clearly by Fig. 30^60^. Combining the result in Fig. 30 with the relation given in [61] that *e*(*t*) *< T*, where *T* is the age of the universe, or maximal propagation time, which leads to the well-known relation between the Hubble parameter and the age of the universe 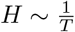^61^.

**Figure 29.**
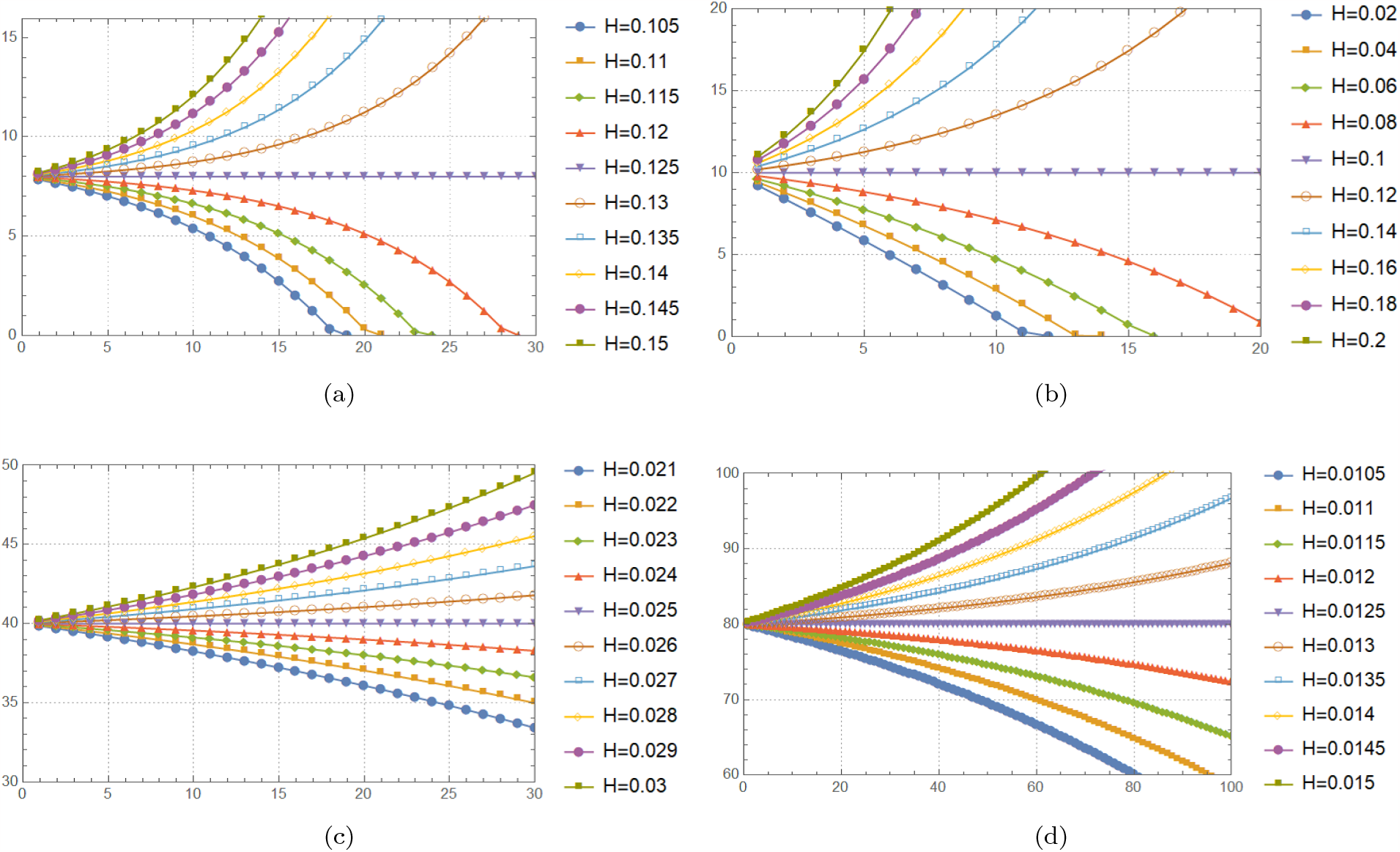
These plots show the expected evolution of light-like trajectories in an expanding space, with *H* = *p*_*d*_ − *p*_*r*_ and *p*_*r*_ = 0, and where space is represented by the vertical axis and time by the horizontal. Natural horizons can be seen to occur at 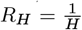

**Figure 30.**
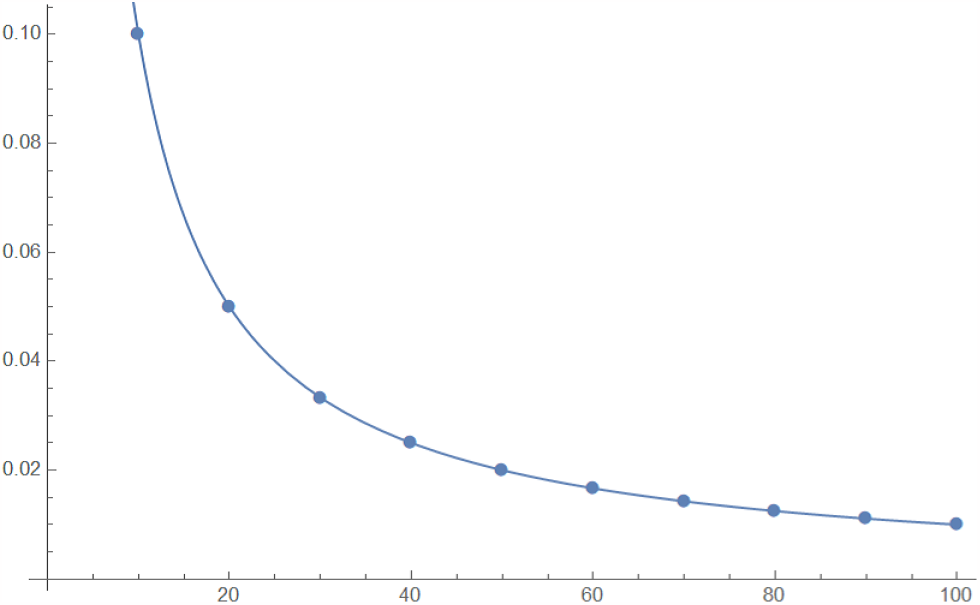
This graph shows the relationship between the constant Hubble parameter *H* (vertical) and the radius of horizon *R* (horizontal). The dots represent values of the Hubble radius calculated through experiment. The line represents the function *H* = 1*/R*_*H*_.

#### Definition 4.1.

The ***scale factor*** in the continuous limit of exponentially expanding SERD network is modelled by *a*(*t*) ∼ *e*^*Ht*^.

#### Lemma 4.2.

*In an exponentially expanding SERD network, at large enough scale, the* ***Hubble constant*** *H satisfies the following relation: H* = *p*_*d*_− *p*_*r*_, *where p*_*d*_ *and p*_*r*_ *are the effective propabilities of duplicaton and reduction respectively*.

*Proof*. The scale factor for an exponentially expanding space is given by definition 4.1. Therefore we can attempt to calculate the total space-time volume spanned out by the expanding IE between two points in global time *t*_0_ and *t*_1_ with *t*_0_ *> t*_1_. Therefore, 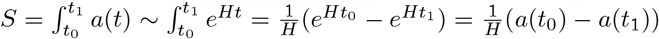. Then given that *δ* − *ρ* = *a*(*t*_0_) − *a*(*t*_1_) where *δ* and *ρ* are the total number of duplications and reductions which occur within the IE in the time interval, and combining with definition 3.9 and 3.10, the result follows. □

#### Theorem 4.3.

*The* ***scale factor base*** *of the SERD network ϕ is equal to e*^*H*^, *where e*^*H*^ = 1 + *H*.

*Proof*. The scale factor increases in increments of time with *δ*(*t*) − *ρ*(*t*) = *a*(*t* + 1) − *a*(*t*), where *δ*(*t*) and *ρ*(*t*) are the number of duplications and reductions which occur in a given TS *t*. Since *a*(*t*) ∼ *e*^*Ht*^ then this equation reduces down to *ρ*(*t*) = *δ*(*t*) + *a*(*t*)(1 − *e*^*H*^). Then at large enough scales, using lemma 4.2 we may say that 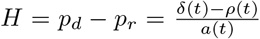. This gives *δ*(*t*) = *Ha*(*t*) + *ρ*(*t*). Then substituting back into the original equation and simplifying, we get *a*(*t*)(*H* + 1 − *e*^*H*^) = 0, and we have the desired result, 1 + *H* = *e*^*H*^.

#### Theorem 4.3

provides a connection between the parameters of the SERD network model and standard cosmological parameters, in this case the *Hubble constant*. We show that it relates to the difference in probability of duplication and reduction at sufficiently large enough scales. It also describes an inflation as a simple powering of a number where we can say *a*(*t*) = *ϕ*^*t*^.

#### Theorem 4.4.

*The expected trajectory of a PSB produced from a split that occurs beyond a COPPs Hubble horizon will never reach the COPP in the growth-only model*.

*Proof*. When splits occur, they generates a set of radially propagating PSBs which follow *light-like* trajectories in the direction of all other PPs in the system, thier propagation speed in the *hidden space* being *1 SE/TS*. So long as we take *p*_*r*_ = 0, given observations of expected *light-like* paths in this space from Fig. 29, and equation 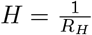, from Fig. 30, we have an effective limit on the region within which *light-like* trajectories will reach the COPP. Since PSBs follow expected *light-like* trajectories any expected trajectory of a PSB occurring outside the radius *R*_*H*_ will never reach the COPP.

This theory is naturally incomplete, namely since we take *H* to be constant and we ignore the effect of SE reductions. Also this will lead to a *static horizon*, and a constant age for the universe, which naturally is not the case. With the inclusion of reductions it is likely that we may see some of these issues rectified^62^. Nevertheless, the expanding SERD network has the capacity to create causally separated regions with no information transfer between them. Fig. 31 demonstrates this effect. Each branch cluser circled in red corresponds to a separate universe, in which information may propagate within. However the expansion of the space keeps these regions of space separate with no information transfer between them. In this context we can imagine separate universes, in an eternally inflating multiverse, as branch clusters of fully connected radially spanning trees.

**Figure 31.**
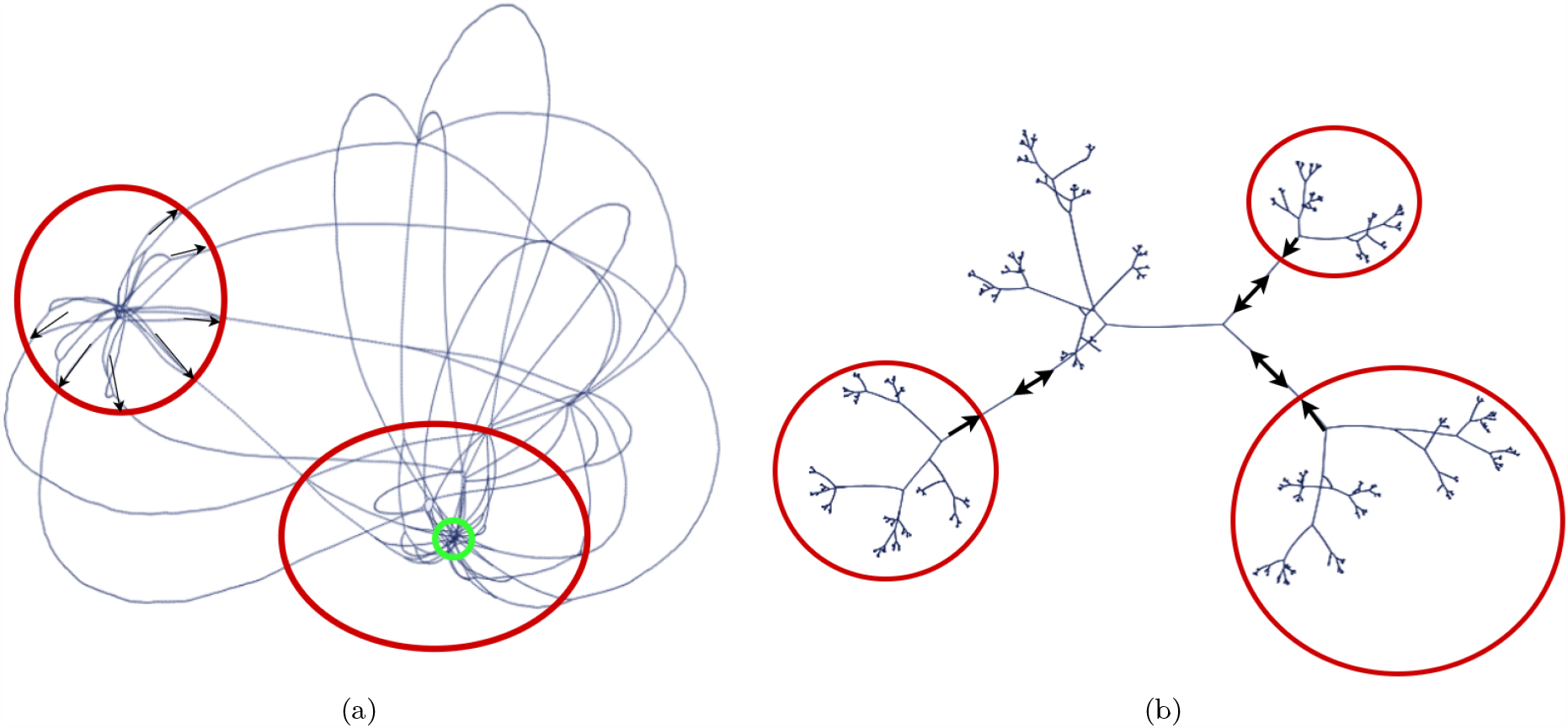
(a) Causally separated regions of the SERD network, with the red circle showing the PSB wavefronts, within which are the PPBCs, and the green circle representing a chosen COPP. (b) Radial subgraph of the SERD network embedded in 2D space with *spring electrical embedding*. The red circle represents a PPBC, the information of the existence of which is has not updated on the COPP.

### 4.5 Correspondence between redshift scale factor relation of FLRW metric and information dispersion in expanding SERD network

When actions occur - in or at the end of an IE - through a TS, the information of that action travels as a *superposing wave-front* or propagating *bit* of information, in both directions along that IE. For SE actions, inside IEs, this results in a number - either a − 1 or a +1 - being added to a number stored on the IG node. These nodes are binary, like a two-way street„ with a left side and a right side, one number that moves ‘*left*’ *a*_*l*_(*t, x*) and the other ‘*right*’ *a*_*r*_(*t, x*). This information propagates along the IE radially in both directions ending up on and updating a PPs effective spacetime metric *e*_*ij*_(*t*)^63^.

Every TS a set of numbers, one for each non-radial IE connected to the PP, which are transferred across the PP as an array to the neighbouring IG of the radial IE connecting the COPP. This means that for a radial IE a constant flow of arrays is passed to the IG neighbouring the bounding PP.

From definitions 3.16 and 3.17 we have a set up where two unit PIPs may superpose due to SE reduction generating a PIP with an effective magnitude of 2. If two PIPs superpose via reductions then the effective magnitudes add together. When superposition of the unit PIPs occurs, it can be thought of as generating an array of concatenated arrays with the effective magnitude being the length of the array of arrays on the IG^64^.

#### 4.5.1 Mathematical correspondences of SERD network expansion and redshift scale factor relation

In this section we shall undergo some mathematical analysis of the information content of the PIP at the moment of observation for a PIP which propagates from a given location in a nonexpanding or stationary space and one which propagates through an exponentially expanding space^65^, modelling both duplications and reductions. By treating the scale factor as proportional to the exponential *a*(*t*) ∼ *e*^*Ht*^, as per the inflationary era and dark energy era, we introduce mathematical models with which to analyse the generation and information depletion of PIPs.

Let us start by considering a PIP emitted from a PP at a time *t*_*e*_ and observed at a time *t*_*o*_, and attempt to model an approximate form for the final information content of the PIP at the moment of observation. Then we will define this for a static space that is not expanding and take the ratio of the two. When an apostrophe is used it implies that the variable in question is defined in the expanding space.

##### Definition 4.2.

*A* ***reduction contraction*** *R*(*t*) *is defined as a number of connected SEs which neighbour the location of a PIP that reduce in the same TS, causing all contained information to superpose and contribute to the overall energy of the PIP*.

Throughout this section we will refer to a *large-scale approximation*. This simply means that the PIP was emitted when the IE size *a*_*e*_ is very large^66^ This will be useful later in our calculation.

##### Theorem 4.5.

*The expected size of a reduction contraction* 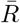 *is equal to the probability of reduction p*_*r*_. *Proof*.

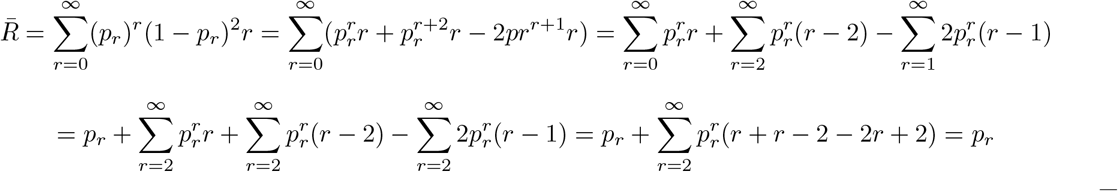

□

##### Definition 4.3.

*The* ***observed information content of a PIP*** *is given by E*_*p*_, *and relates to the number of unit PIPs emitted from bounding PPs contained within the observed PIP due to superposition via SE reduction, and it may be calculated using the following eq. 3*^67^.

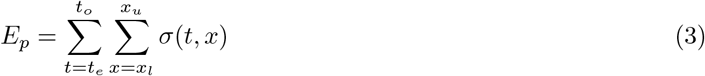

*Where t*_*e*_ *and t*_*o*_ *are the times for emission and observation of the PIP respectively, with τ* = *t*_*o*_ − *t*_*e*_. 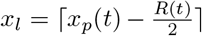 *and* 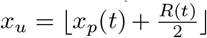. *Here R*(*t*) *is the size of a reduction contraction at a time t and σ*(*t, x*) *is the propagating information content of the IG x at time t from a non-observing bounding PP*..

In this case there is no distinction between *left* and *right* since we are only interested in one direction - that is the direction inward towards the COPP or *observer*.

##### Definition 4.4.

*The* ***average IE information density*** *in the* ***close neighbourhood of IG*** *x is given by ϵ*(*t, x*) *at time t*.

##### Definition 4.5.

*The* ***average local IE information density*** *associated with a* ***close neighbourhood of a PIP*** *p is given by ϵ*_*p*_(*t*) *at time t*.

When we average the value of the observed information we may replace the specific information quantities *σ*(*t, x*) with an average information density at and in the neighbourhood of the PIP *ϵ*_*p*_(*t*). Since *a*(*t*) is large and the function *ϵ*(*t, x*) is smooth and slowly changing over *x* and *t*, as is demonstrated by Figs. 32 and 33, we may make the approximation given by equation 4.

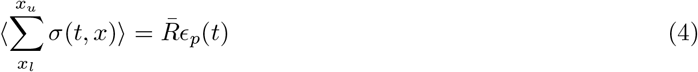

Where 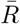 is the average size of the reduction contraction which is equal to *p*_*r*_. This paramater could be both time and scale dependent; however, we explore the special case where it is neighther^68^. *x* is the location of the photon or PIP at time t.

Since 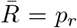, we are only varying the function *ϵ*(*t, x*) over a small distance so we can approximate the integral over space as 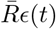. In the *large-scale approximation* we may take the continuum limit^69^.

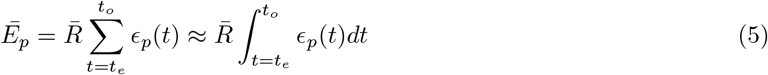

##### Definition 4.6.

*The* ***average PIP wavelength*** *of PIP p is givem by λ*_*p*_(*t*) *and represents the average distance between* ***bits*** *or* ***information packets*** *at the location of the PIP at time t*.

##### Theorem 4.6.

*Given the assumptions of large-scale continuity, exponential expansion and a Hubble parameter constant in time* 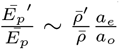.

*Proof*. By definition the *average local IE information density* and *average PIP wavelength* are related by equation 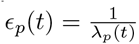, this then leads to the equation for the observed PIP energy 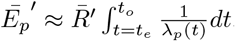. As described at the start of this section, unit PIPs are emitted every TS and move inward toward the COPP. Reductions cause the superposition of PIPs, increasing the *effective magnitude* of the inward propagating PIPs. At the same time duplications - of which there are *a*_*o*_ − *a*_*e*_ more of overall - cause the PIPs to become sparser as they propagate inward as demonstrated by Fig 32 for *growth only* and Figs. 5 and 33 for the inclusion of reductions. Reductions occur *homogeneously* though the IE on average and so when considering the value of *λ*_*p*_(*t*) we can consider the growth only model since the reductions cancel out. Since unit PIPs are emitted from the bounding PP every TS *λ*_*p*_(*t*_*e*_) = 1, then as the system scales the distance between these two points grows along with the scale factor leading to 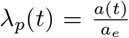. Therefore we can write 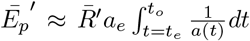. Treating the scale factor as an exponential *e*^*Ht*^ ^70^ and making use of theorem 4.5 and definition 3.9 leads to 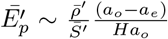. Since we are in an exponentially expanding space we can make the following definition for 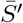, the average number of spacetime events in the time interval, 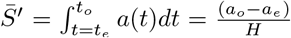. Combining these gives 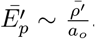. Now we consider the stationary frame. Since there is no overall growth and the IE is static on average we can say *ϵ*(*t, x*) = 1 ∀*x, t* ∈ ℕ. This means that we can write 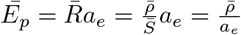. Since in the static case the update time is simply equal to the scale at time of emission *a*_*e*_, and *S* is equal to the distance times the update time which in this case is equal to *a*_*e*_. Bringing all this together leads to the final result^71^.

#### 4.5.2 Computational results showing satisfaction of redshift scale factor relation in SERD model

As discussed in *subsection 3.5*, the FLRW metric leads to a relation between the ratio of the scale factor at emission and observation time, *a*(*t*_*e*_) and *a*(*t*_*o*_) respectively. We provide computational results of simulated information transfer radially in the expanding 1D single IE SERD network and show, using the methods outlined in *Subsection 3.5.2* correspondence to cosmological red-shift predicted by the FLRW metric.

*Subsection 3.5.2* gives a description of the implementation method that we used to computationally test this relation in the SERD network model. In this subsection a comparison is drawn between the redshift, or energy/information loss, of photons through expanding space, and the loss of information of PIPs when propagating through an expanding space in the SERD network.

We make an assumption, similar to that in cosmology, that we may extrapolate on the observed spatial expansion beyond the observed horizon. That is that space expands on average at the same rate everywhere at a given instant, or that there is no spatial dependence of the cosmological constant. Therefore the scale factor *a*(*t*) is the instantaneous number of SEs in the hidden state of the IE.

At every TS, a PP sends out a *unit PIP*, encoding the information that has reached the bounding PP, in that TS, through other connected IEs, and then ir propagate inward towards the central PP. This can be imagined as a steady stream of bits (1s) entering one end of the radial IE each TS, superposing into the

PIPs with specific PIP magnitudes due to reductions and then updating on the COPP, informing the COPP of the action information that has reached the bounding PP. This can be seen in Figs. 33, showing how the PIPs average out over the sample of Monte-Carlo runs into an average energy density.

Fig. 34 shows how the average *effective PIP magnitude* decreases as we increase the rate of expansion of the system. This shows direct evidence of expansion induced redshift but does not demonstrate direct correspondence to the redshift scale factor relation of the FLRW metric. Fig. 35 plots the values of ^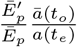^ for specific values of *a*(*t*_*e*_), with *Ē*_*p*_ representing the average PIP magnitude or energy of the PIP. In order for the redshift scale factor relation to be satisfied this value must approach one at the correct scales and with sufficient samples, as is observed close to the correct scaling in the results presented in Fig. 35.

**Figure 32.**
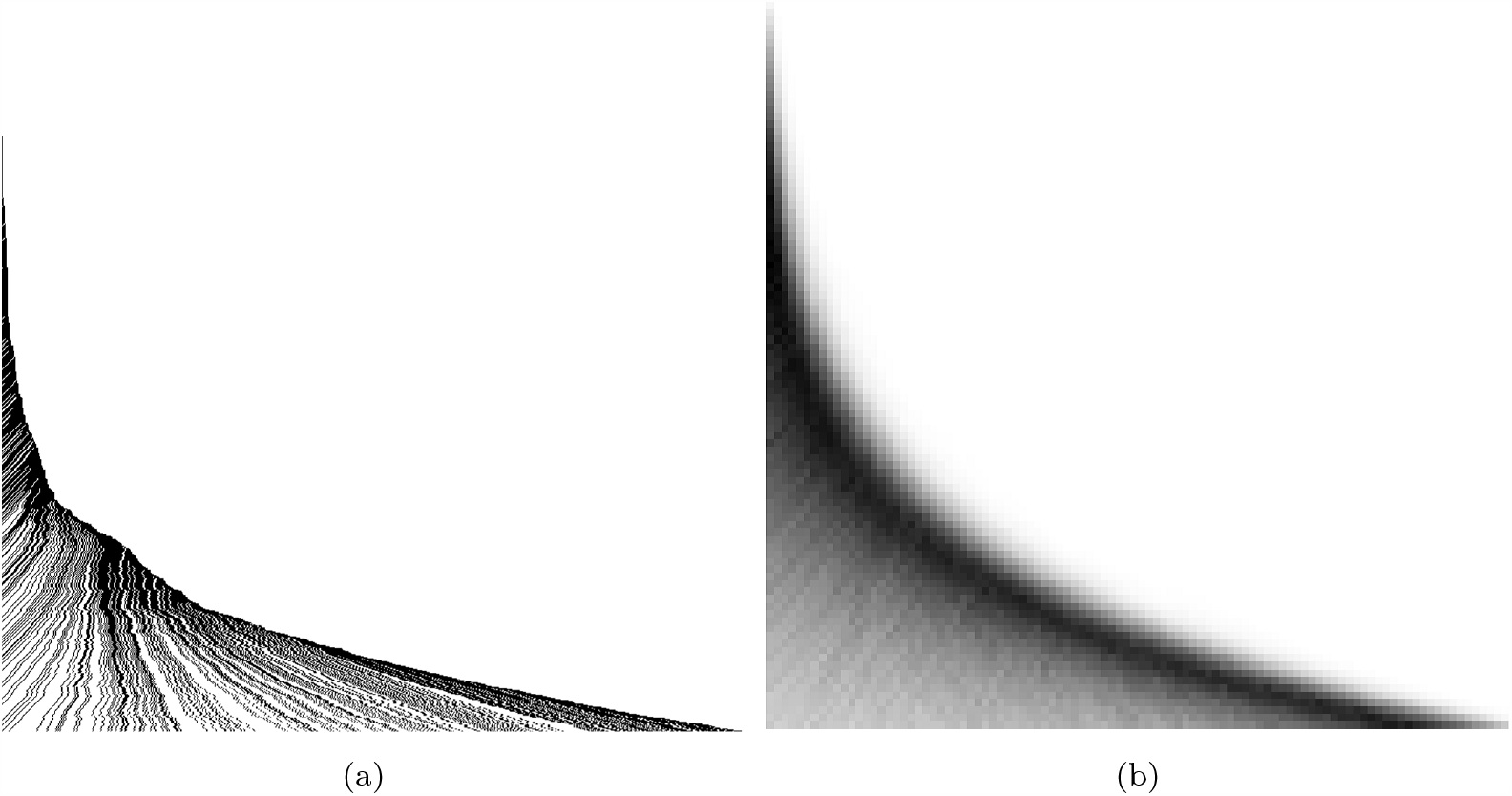
(a) Temporal evolution (vertical downwards) of the *propagating information* due to PIPs for the evolution of one branch of an exponentially growing 1D IE (horizontal) within the SERD model not including reductions. Bits of information are produced from an enclosing PP every TS and then dispersed across the IE as the IE expands and the information travels across it. (b) Temporal evolution (vertical downwards) of the *average information density* for all branches that are constrained to have a global scale factor of between 90 and 110 for 100 TSs of a growing 1D IE (horizontal) within the SERD model not including reductions. The average energy density decreases as the scale of the system increases. The expansion of space dilutes the spatial information density.

**Figure 33.**
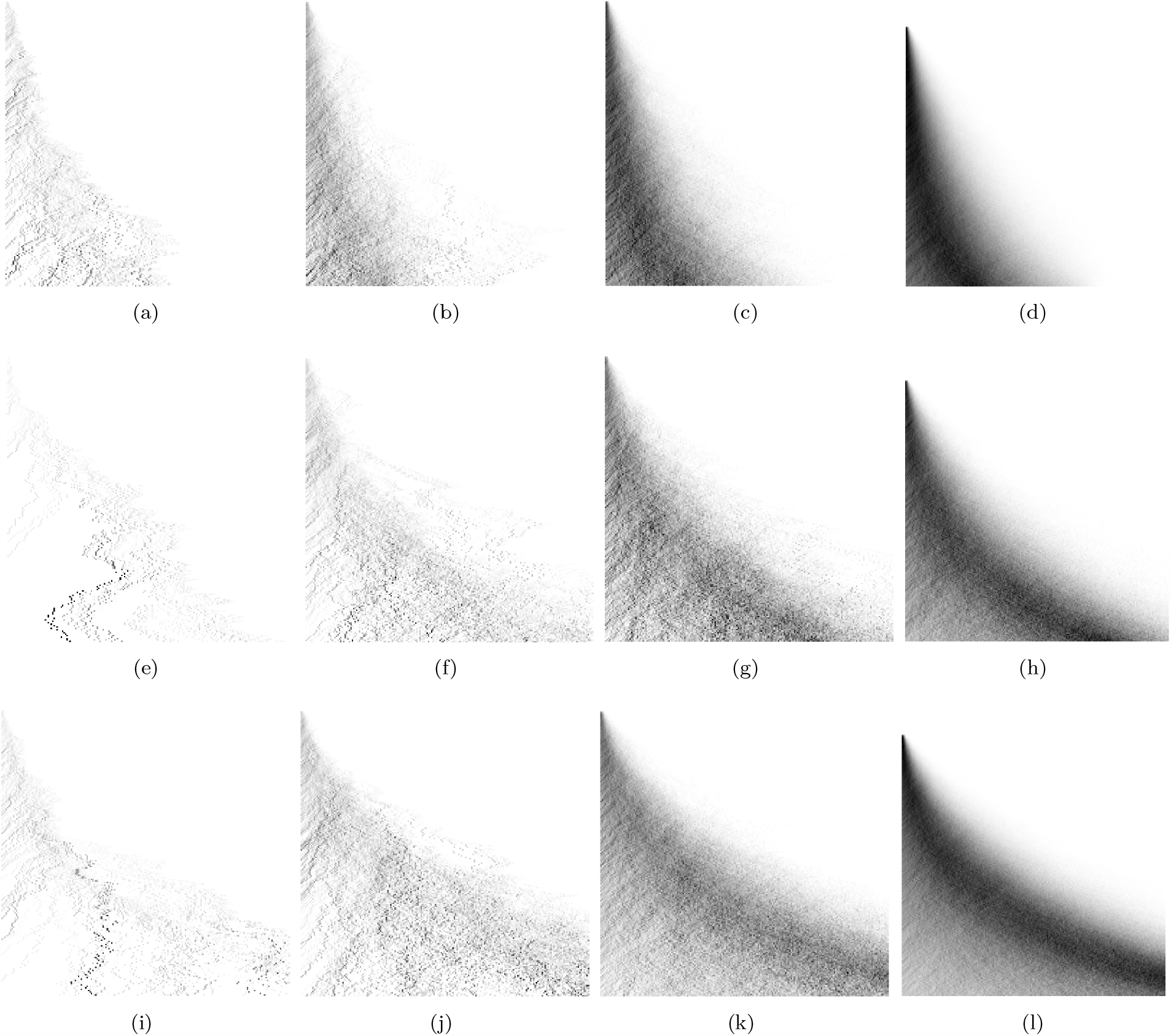
These figures show the space (horizontal) time (vertical) diagrams for an ensemble of sample IEs evolving for 200TS. The darker spots are the higher-magnitude PIPs propagating from right to left as the system inflates exponentially. Rows (a) to (d) scaled to between 50 and 150 SE, (e) to (h) 150 to 250 and (i) to (l) 250 to 350, with the number of iterations starting at 100 and going up in powers of 10 from left to right.

**Figure 34.**
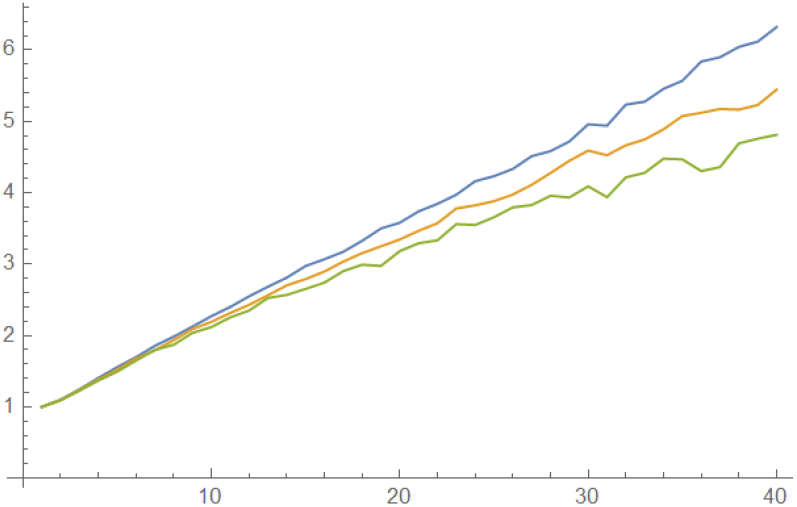
This plot shows average PIP magnitude (vertical) vs time (horizontal) for a single IE with varying expansions. Starting from a single SE state and growing to 40SE (blue), 80SE (yellow) and 160SE (green) respectively. The redshift is clearly visible, however due to the small scale of the system implemented this redshift is unlikely to correspond directly to the *redshift-scale-factor relation* of the FLRW metric. Nevertheless, it does clearly demonstrate that PIP internal energy is decreased due to expansion.

**Figure 35.**
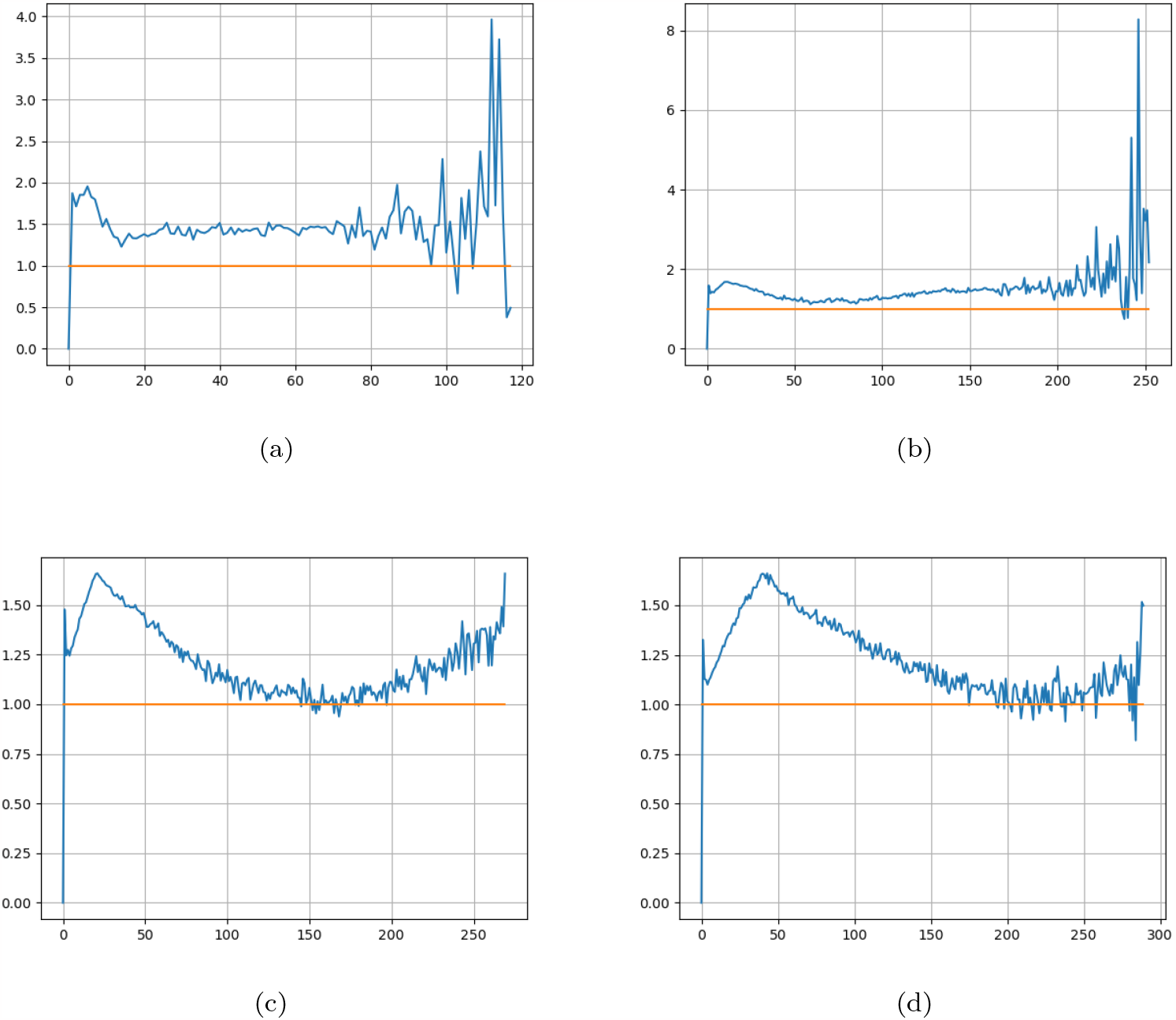
Graph showing the value of 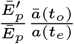 against *a*(*t*). (a) implements a first tier for 100TS for 10^8^ sample runs giving 2283 inputs within range 40-60SE for second tier of 100TS with 10^8^ sample runs. (b) implements a first tier for 200TS for 10^8^ sample runs giving 1351 inputs within range 80-120SE for second run of 200TS with 10^8^ sample runs. (c) implements a first tier for 400TS for 10^7^ sample runs giving 738 inputs within range 160-240SE for second tier of 400TS with 10^7^ sample runs. (d) implements a first tier for 800TS for 10^8^ sample runs giving 314 input states for the second tier of 800TS for 10^8^ sample runs. In each case sample PIPs were observed within a range of 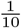times the second run time duration and the final state and averaged.

## 5 Discussion

At the heart of this paper is the idea that the observed connections between large-scale cosmological structure and pattern formation in nature seen in [6] are consequences of the underlying atomized space-time network, and that the branchial dendritic structures which occur in nature provide us a clue as to what this underlying structures may be. Further, that life follows these patterns as they from pathways of least energy cost for both matter and energy flow [89]^72^.

We investigated a specific discrete spacetime model, the *SERD network model* introduced in [59, 60, 61] due to its emergent hypothesised similarities to mycelial structure and evidenced physical comparisons. **A key point to make here is that the SERD model was never designed to generate mycelial structure, but emerged out of more fundamental considerations** [61]. Investigations into this model took place long before any connection to mycelium, or patterns in early evolving life, was made. This is what makes this model of significant interest to the field, as it forms a bridge between three main areas of research; evolution of network structures in early life, cosmology and discrete spacetime.

The SERD model is an evolving hypergraph^73^, where the edges are elements of space, the hyperedges are point particles, elements of matter, and the nodes are the gaps between the elements where propagating information is stored. It evolves via a combination of non-deterministic (element actions)^74^ and deterministic (information transfer)^75^ update operations. Information of actions propagate long edges at a rate of one edge per time step and are updated on the point particles that behave as the observers in the SERD network.

We draw connections between the elements of this model and elements in a mycelial network and imply, with the aid of diagrams and computer simulations, how the morphology of mycelium can be realised in the SERD network.

Here we introduced two main implementations of the SERD network model, the *growth-only model*, and the *single-interaction-edge model*.

The *growth-only model* is an evolving network implementation of the SERD model which excludes reductions^76^. The results showed the first images of one branch of the SERD network’s evolution^77^ with emergent spanning-tree radial topology, observed non-trivial connections between hyperedges and interactions of propagating branches. A toy model of information transport inwards along these radial trees was also introduced. The impediment and dispersion of propagating information due to the exponential growth of the space of the SERD network model was also observed^78^. Connections between the exponential expansion of space in the SERD network and exponential expansion of space in ΛCDM model was made. The Hubble parameter was fixed as constant and defined in the context of the SERD model. The exponential expansion of the space and constant information propagation speed generate horizons, beyond which information cannot reach an observer, resulting in causally separated regions of space and *pocket universes*^79^. An inverse relationship between the Hubble parameter and the radial size of this horizon was observed.

The single-interaction-edge model allowed for us to investigate the effect of reductions and duplications on propagating information packets emitted from hyperedges. Information which has reached a hyperedge is emitted as a unit of propagating information at every time step, sent out as a signal to all other hyperedges in the network^80^. As these bits of information transmit the information of update operations through the network both the dilution of information - via *duplications* - and the superposition of information - via *reductions* - are observed. This results in a specific magnitude of information reaching the observing hyperedge which it updates on at a given time. We make the connection that these propagating information packets behave like photons, with the energy of the photon being defined by the magnitude of the information on it at the time of observation.

Some mathematical analysis was done on the evolution of propagating information in this model given the assumption of continuous exponential growth, however this stopped short of direct correspondence with cosmological models with a factor of the ratio of total number of reductions between the stationary and expanding frames^81^.

We demonstrate through computational implementation of the single-interaction-edge model that as space expands in the network, it has the effect of reducing the overall magnitude of the propagating information packets. **Most importantly, we show strong correspondence between this effect of expansion-induced information reduction of propagating information packets and the cosmological *redshift-scale-factor relation* of the** Λ**CDM model presented in Fig. 35**.. This strengthens the argument that these propagating information packets behave as photons do as they propagate through space.

The fact that this model shows direct cosmological, as well as a number of other documented physical [59, 60, 61], comparisons and that it hints at mycelial topology justifies further research into this system as both a possible underlying spacetime and as a model for growth and information transfer in mycelium and other biological networks. It may also help in deepening our understanding of the connection between early complex life forms and large-scale structure in the univserse.

We will soon have further results to release relating to the inclusion of reductions in the SERD network, implementing smoothy embedded evolutions of the network as well as further mathematical analysis. Including reductions in the code will not only result in observation of lateral branching and anastomosis, but it will also open the door to exploring the compactified spaces of particles, the superposition of propagating branches and the observation of the repression of the growth of these branches^82^ to name just a few potential applications.

The SERD model is a highly dynamic information-processing machine which emerges from simple rules and generates both spanning-tree radial structure and physical comparisons. It has many features at varying levels of complexity. This paper has progressed our understanding of this model, making further physical and biological comparisons and presenting the emergent biologically similar networks it generates. Further research into this model will probe the deeper layers of emergent complexity it generates.

## 6. Acknowledgement

The research has been conducted under the framework of the FUNGATERIA (www.fungateria.eu) project, which has received funding from the European Union’s HORIZON-EIC-2021-PATHFINDER CHALLENGES programme under grant agreement No. 101071145. It is co-funded by the UK Research and Innovation grant No. 10048406. We are grateful to Ella Schunselaar, Han A.B. Wösten, Koen Herman and Robert Jan Bleichrodt for providing experimental results and videos on growing mycelium of *S. commune*. We are also greatful to Leon Conrad for his help and support during this papers final stages.

## Footnotes

1 Some examples of these difficulties; combining the two theories in extreme conditions (such as those in and around a black hole, the earliest moments of the big bang, the centre of a neutron star), defining the quantum behaviour of particles as world lines on a continuous spacetime manifold, the hierarchy problem, the problem of non-renormalizability.

2 Patterns of energy discharge and optimized energy flow.

3 This paper provides a motivation for hypothesising a correspondence between this model and mycelium networks via diagramatic demonstration of the mechanisms to realise apical and lateral branching, fusion, mitosis and cell death (*section 3.3*). However the formal network implementation is still incomplete and so direct correspondence is not yet observed in the results.

4 Sometimes referred to as filamentous, or higher, fungi.

5 Also referred to as an inoculum.

6 The first stages of the mycelial hyphae.

7 Depending on the species of mycelial fungi investigated, different amounts of apical and sub-apical branching may occur. Most appears to be sub-apical, resulting from an effect refered to as apical dominance, where branching events tend to take place a given distance away from the extending hyphal apex.

8 The opposite affect may occur since increasing the minimum number of junctions between source and sink will have the effect of decreasing the flow rate due to resistance in the network [75].

9 Compared to processing speeds of modern computers.

10 This may make mycelium networks a useful substrate to work with when producing responsive biomaterials and unconventional computers for specific tasks.

11 The *Bejan constructal law* states that for a flow system to persist in time it must evolve freely such that it provides greater access to its currents. In this specific case a flow system may be a living mycelium, but it may also refer to any living organism that persists in a state of input output energy flow with the environment. Currents is a general term but refers to the currents that nourish the organism, such as flow of nutrients, glucose and energy singals, but could also refer to blood flow, oxygen or food. This cans be generalised further to flows of energy and matter.

12 Hyperedges can be connected to any number of nodes.

13 Edges can only connect two nodes.

14 The SERD network updates via a combination of non-deterministic and deterministic update operations and so can be considered a *mutliway system*.

15 In the computation of the SERD network these labels are numerical-symbolic interchangable, which allows for the label to act as both an identifier and an address.

16 The mathematical structure of the propagating information on an IG may vary depending on the level of detail of the SERD network that is implemented. This maybe a PIP object or a PSB object, but this may vary depending on the implementation. In some implementations the information for PSBs is stored in the IG set where as in other implementations the information for PSBs is stored in a separate PSB set.

17 In this paper we have excluded this action from the network evolutions (not the one dimensional evolutions) due to implementation challenges relating to superposing PSBs and time constraints.

18 The merge process is not investigated in this paper, but it will be included in more comprehensive simulations in future papers.

19 This number of PPs that a PP is aware of is exactly equal to the number of IGs connected to it.

20 This may be done via addition, array addition, or set concatenation, depending on the level of detail that is being probed in the network,

21 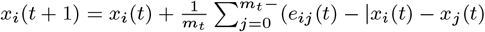

22 Further evidence of this maybe found on the mathematica notebook found on the following github page given in the additional references section.

23 Since the embedding always starts from a unique initial state and then depends only on earlier embeddings of the evolution we may still refer to the SERD network as *background independent*.

24 For PSBs, in contrast to the PIPs, the information is stored in the connections between elements, or the topology of the network, which information in PIPs is stored internally as an information carrying object such as an integer or array.

25 Figs.**??**,20,21,23,24 and 31.

26 Confluence of multidirectional PIP and PSB propagation within a full SERD model system representation is not included in this paper and may be the subject of further consideration in later papers. However evidence of confluence of overlapping PSBs is demonstrated by Fig8.

27 Additonal evidence of physical comparisons not included here are; Newtons laws under *time iterated embedding*, sum over histories, compactified PP spaces, constancy of speed of light in all inertial reference frames under *time iterated embedding* and elementary measurement.

28 They superpose on one another with no limit to the amount of information density in space (bosons), PPs however are separated in space and so naturally exclude one another from the same regions of space (fermions).

29 A *growth-only model refers to a version of the SERD network model that only implements PP splits and SE duplications but not reductions*.

30 Sub-apical lateral branching is a process that requires the implementation of reductions to observe and will not be observed in the *growth only model* explored in this paper.

31 This also means that in the SERD model the ‘seed’ of the hyphae branch is always created at the apex but remains dormant until it passes the scale threshold. There may also be a relation to *apical dominance*.

32 These PPs then form a *Connected PP Network* (CPPN).

33 Anastamosis is a process that requires the implementation of reductions to observe and will not be observed in the *growth only model* explored in this paper.

34 When considering the observed size of the system [61] is constrained by the overall age, where as the hidden state may expand exponentially eternally.

35 If a breakage was to occur due to SE reduction, then what would define the space of the separation? The connectedness of the SERD network is one of its most fundamental features, and although regions maybe separated via inflation induced horizons [61]. If reductions caused breakages in the IEs, then the SERD network would almost immediately become a series of disconnected SEs, with no meaningful structure or dynamics.

36 It could be speculated that this natural attraction to large IEs of PPs with reduced sub-IEs to a PSB on the IE results in the threads of the biologically similar network of dark matter observed in our universe at large scales.

37 The average observed growth rate of IEs is constrained to below the speed of light in the physical comparison of the SERD network and is 1 SE per TS as demonstrated by results in [61].

38 This operation of update of information could be described as more fundamental than photons, relating to any matter interaction of a force carrying boson.

39 Monte-Carlo simulations allow for testing expectation values along large time lengths. For testing cosmological redshift, since we are only considering one photon over one dimension, it is suitable. However it is possible that certain, most likely quantum mechanical, physical comparisons are lost due to the requirement of the effective observation of each hidden state at every moment in time through each run. This does not allow for a more efficient algorithm that only stores information of what is observed rather than what is hidden.

40 A *sample run* may be considered as a single branch in a multiway system or number of loops in the give tier of the Monte-Carlo simulation.

41 This magnitude is directly proportional to the information content of the PIP.

42 Therefore the input state of the first tier is given by *α*(0) = (0, 1)

43 These mechanisms are implemented using the .insert() nd .pop() operations in python and are defined as (…, *α*(*t, x*), …) → (…, *α*(*t, x*), 0, …) for duplication and (…, *α*(*t, x*), *α*(*t, x* + 1), …) → (…, *α*(*t, x*) + *α*(*t, x* + 1), …).

44 A suitable length of time *t*1 is required to allow for a suitable sample size to grow to *s*1 and also to stabilise. The difference between the dilated time *t*′ and proper time *t* gives a measure of how close the system is to *static equilibrium. Static equilibrium* is the theoretical most likely thermodynamic limit of the internal information distribution of the IE.

45 For example, one may implement a single IE with duplications and reductions but no information propagation, then one may implement a model with splitting but no reductions or information propagation. There are ways to investigate specific aspects of the network without requiring the development of code to implement every feature of the network model.

46 This will be the topic of a follow-up paper

47 Albeit somewhat dampened by the exclusion of SE reductions

48 Another way to define this is that *p*_*r*_ → 0.

49 This code is simpler than one which includes reductions since it does not require deleting elements and IGs, updating SE neighbours in PSBs and superposing information on IGs. The process of coding the SERD network is incremental and once we write a full code, including reductions, we shall observe many more detailed behaviours. This will likely be the topic of a follow up paper.

50 This is directly due to the fact that the lack of reductions prevents IEs from decreasing in size and so once they get large enough that ne IEs, generated from splits, that start with a size of 1 SE will never overtake the IE scale between PP clusters.

51 Early stage of a hyphae.

52 These interaction patterns can be observed in Fig. 25, 26 and 27.

53 Normally 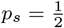 since all actions in the SERD network model are treated as being equally likely.

54 In the mycelium hyphae propagate for longer before branching, whereas in the SERD network apical branching is occurring at pretty much a constant rate.

55 In a hypothesised mycelium comparison between spacetime and mycelium one can imagine that the mycelium network is centred around one observer within a specific observable boundary. Then when particles fall past a horizon, either cosmological or a black hole, this is like a spore travelling to a new location to generate a completely new mycelium network that is causally separated from its parent mycelium. In this way, different observable universes are thier own mycelium networks, seeded with the PP at the hyphal tip or IE that has crossed over the horizon.

56 It is important to note that these cycles only occur when two PSBs travelling in opposite directions cross paths and so do not happen for the radial sub-graph of the COPP if the COPP has its splits repressed.

57 Note that we are observing a reduced implementation of the SERD network model rules.

58 This may have some interesting correspondences with quantum phenomena since this may allow for individual photon self-interactions. If PIPs must ‘choose’ which IE to propagate along at a PSB branch this may also correspond to the *inverse square law* of radiation dispersion. These two behaviours may be mutually exclusive and depend on whether PIPs follow all branches, or just one, at a PSB branch.

59 A more quantifiable correlation with mycelium and slime mould is now possible and will be investigated in more detail in a future paper.

60 The Hubble radius being the distance beyond which information may never reach an observer.

61 This is a short hand approximation of the universe’s age that assumes the Hubble constant is constant in time and that the universe is expanding at a rate directly proportional to t. This is of course not the case since the Hubble constant is not constant in time and the universe does not expand at a constant rate. Also, this would lead to a single value of the age of the universe which of course would not be true. Nevertheless, this correspondence does demonstrate some connection between how horizons form in the SERD network, and in modern cosmological models.

62 If reductions are included, there is always some non-zero possibility of a PSB to crossing a horizon that it would not be able to in the *growth-only model*, this probability drops to close to zero if the system grows and the PSB is located futher from the horizon, but it leaves open the possibility of an ever-growing horizon, such as the one derived in [61].

63 In this paper, a full bi-directional model, with PIP self interactions is not implemented since we are focussing on the radial expansion properties of the model and connections with red-shft scale factor relation. As a result a more comprehensive simulation of all SERD model rules and behaviours is not yet explored. Perhaps PIP branching and looping behave like self interacting wavefunctions when specific PIP path is not observed.

64 This specific implementation is not modelled in this paper instead we are interested in the effect of 1D-scaling and how this scaling affects the observed information content (or energy) of the PIPs.

65 One major assumption in this is that the growing SERD network can be modelled by continuous exponential growth.

66 Precisely it may make sence to say that *a*_*e*_ reaches sizes of 10^45^ since the universe is now about 10^60^ planck lengths accross and at this point this is the most natural scaling. However in computational results (Fig. 4.5.2) we show that we reach physical correspondence at much smaller scales.

67 This equation ignores edge effects since *p*_*r*_ is so much smaller than the scale of the IE.

68 It will be shown later how this relates to keeping the Hubble parameter constant. We may also explore how time and scale dependences effect the resulting derivation if time allows.

69 Throughout the following derivation we shall be making a *large-scale approximation*.

70 Given that the fundermental definition of the Hubble perameter is 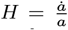 it is natural to think of this as the average amount of expansion one unit of space undergoes in one TS, taking large-scale averages, we can see that in the SERD network model, keeping H constant, that 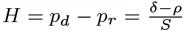

71 This is a preliminary result which may be amended with constraints to the Hubble parameter on the average total number of reductions 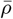 and light-like paths, and considerations of the suitability of the assumptions made. It does not match the redshift-scale-factor relation, however, it introduces some important concepts with which a more formal derivation will be performed in the future and so was important to include it in this paper. We shall see that the results in the next subsection provide a strong proof that the redshift-scale-factor relation is indeed satisfied in the SERD network model.

72 One could use the analogy of iron filings showing the underlying magnetic field structure by falling into local potential wells, or tracing an underlying texture with a crayon.

73 The hypergraph evolves as a multiway system.

74 SE *duplication* and *reduction*, and PP *splitting* and *merging*.

75 PIP and PSB propagation.

76 Including reductions in the code resulted in some implementation challenges relating to accessing data stored in PSBs so time was diverted into solidifying the results from the code already written. Implementation of reductions duplications and splitting will generate lateral branching and anastomosis and will be the subject of our next paper once the coding challenges have been overcome.

77 Reductions cause fluctuation in the scale of the system, and implementation of which will generate far more dynamic behaviour and changing structure in the SERD network

78 It is important to remember that space may expand exponentially in the SERD model, but - as shown in [61] - the observed scale is constrained by the speed of light, as with our own universe.

79 The inclusions of reductions would mean that these horizons are not smooth, but rather rough, with reduction-induced fluctuations of space allowing information and particles in the neighbourhood of the horizon to pass through it in both directions.

80 This information will only reach particles which lie within or are close to the Hubble horizon of the particle which emmitted it.

81 Dispite falling short of direct mathematical correspondence some important definitions were made and may provide a foundation within which to make a more rigerous derivation of the connection between these two models. A number of assumptions were made in this derivation such as constant Hubble parameter and spatial continuity, which on closer exmination may impact the final derived result.

82 IEs or sub-IEs

